# Single-cell DNA sequencing reveals pervasive positive selection throughout preleukemic evolution

**DOI:** 10.1101/2023.12.15.571872

**Authors:** Gladys Poon, Aditi Vedi, Mathijs Sanders, Elisa Laurenti, Peter Valk, Jamie R. Blundell

## Abstract

The representation of driver mutations in preleukemic haematopoietic stem cells (pHSCs) provides a window into the somatic evolution that precedes Acute Myeloid Leukemia (AML). Here, we isolate pHSCs from the bone marrow of 16 patients diagnosed with AML and perform single-cell DNA sequencing on thousands of cells to reconstruct phylogenetic trees of the major driver clones in each patient. We develop a computational framework that can infer levels of positive selection operating during preleukemic evolution from the statistical properties of these phylogenetic trees. Combining these data with 67 previously published phylogenetic trees, we find that the highly variable structures of preleukemic trees emerge naturally from a simple model of somatic evolution in which there is pervasive positive selection acting throughout the disease trajectory. We infer that selective advantages of preleukemic clones are typically in the range of 9%-24% per year, but vary considerably between individuals. At these level of positive selection, we show that the identification of early multiple-mutant clones identifies individuals at risk of future AML.

## Introduction

Large cancer genome sequencing efforts over the last 20 years have revealed that most cancers harbour multiple clonal driver mutations at diagnosis ^1–3^. These mutations are sometimes acquired decades before the emergence of the first malignant cell ^4,5^. How the full complement of driver mutations is acquired in a single cell over a human lifespan, however, is not fully understood. Early multi-hit theories of cancer considering mutation acquisition as the key factor seemingly capture the age-incidence relationships across a range of cancers ^6^. However, using estimates of somatic mutation rates ^7^ and stem cell numbers ^8,9^ these theories struggle to provide plausible estimates for overall cancer incidence ^10^.

Work over the last decade using deep bulk sequencing has shown that there is positive selection acting on mutations in cancer-associated genes across a range of healthy tissues ^9,11–21^. This suggests that positive selection may act throughout the entire multi-hit trajectory of cancer. However, because of the inherent challenges in measuring precancerous evolution, the predictions of this conceptual idea have not been quantitatively tested.

Single-cell sequencing from multiple cells in a population can provide a quantitative picture of the evolutionary history of the population by using somatic mutations as unique lineage markers for building phylogenetic trees ^5,22–24^. The haematopoietic system provides an ideal model for using single-cell approaches as haematopoietic stem cells (HSCs) can be easily isolated and sequenced. These approaches have revealed that large numbers of HSCs maintain the blood in healthy individuals ^8,24^, that the driver events of certain blood cancers can be acquired in utero ^5^ and that there is extensive clonal heterogeneity in acute myeloid leukaemia (AML) ^25–28^. An innovative approach to studying pre-cancerous evolution is through the isolation of ostensibly healthy preleukemic HSCs (pHSCs) from AML patients at diagnosis. By coupling this idea with single-cell sequencing the step-wise nature of preleukemic evolution and how preleukemic clones can seed relapse post therapy has been previously demonstrated ^29^. However, the quantitative information encoded in the preleukemic phylogenetic trees have not been fully exploited to understand how levels of positive selection operate during preleukemic evolution. Here we hypothesize that the statistical properties of phylogenetic trees across many AML patients could reveal levels of positive selection operating during preleukemic evolution and how those levels vary among individuals.

## Results

### Single-cell phylogenies from preleukemic HSCs

To construct phylogenetic trees from preleukemic HSCs (pHSCs) we obtained bone marrow aspirate samples from 16 AML patients collected at diagnosis from the multicenter HOVON-SAKK clinical trials (see ‘Sample selection’ in Supplementary note 1). To minimize inter-patient variation we selected samples with a genetically restricted AML genotype (*DNMT3A*^mut^/*NPM1*^mut^). Preleukemic HSCs were isolated by FACS sorting (Figure 1A, Supplementary note 1). We then performed single-cell DNA sequencing of these pHSCs targeting 45 myeloid genes using the Tapestri platform (Mission Bio). Combining this data with a high-confidence set of driver mutations identified by bulk sequencing of each bone marrow aspirate sample, we were able to genotype a total of 4013 HSCs (range 7 − 988 per sample) for the reconstruction of phylogenetic trees. Individual cells from single-cell sequencing were then assigned to clones based on the presence or absence of these driver mutations, forming a phylogenetic tree based on driver mutations for each sample (Supplementary note 1).

**Fig. 1.**
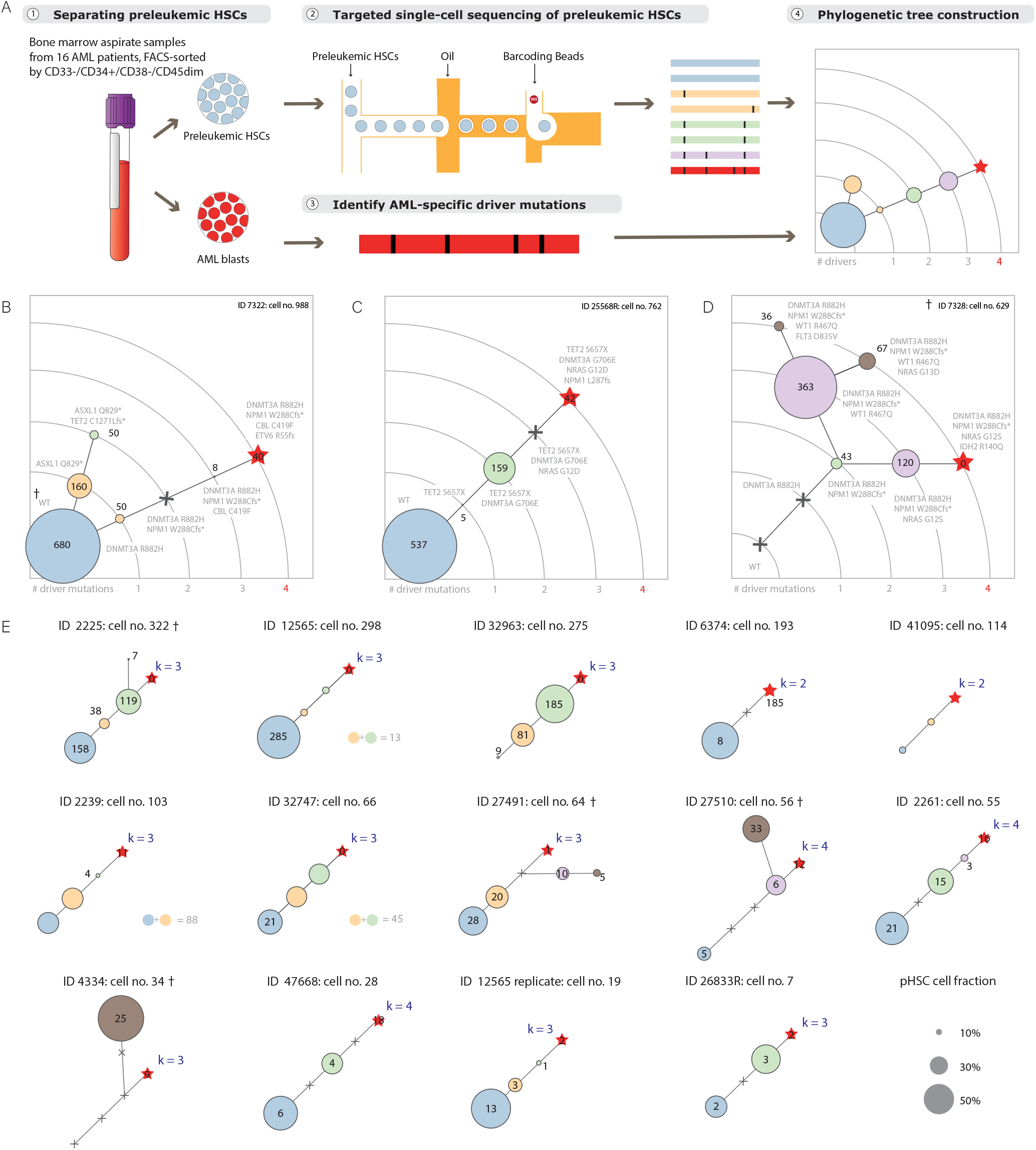
Construction of phylogenetic trees from preleukemic haematopoietic stem cells from AML patients at diagnosis. **(A)** Preleukemic haematopoietic stem cells were sorted from bone marrow aspirate samples acquired from AML patients at diagnosis. The sorted samples were then passed onto the single-cell DNA sequencing platform for genotype information on a targeted panel of 45 commonly mutated myeloid genes. Variant information at single-cell level combined with bulk sequencing information of the same samples before sorting enables the assignment of individual cells to their respective clonal genotype. Phylogenetic trees were constructed for each patient based on the clonal architecture found in the preleukemic HSC population. **(B-D)** Three 4-hit phylogenetic trees from three patients are shown. Clones are denoted by circles and ordered on ascending arcs based on number of drivers detected. The number of cells assigned to each clone is labelled on the circle and replaced by a grey cross if zero. The genotype of a clone is also labelled and represented by a red star if it belongs to the blast genotype. **(E)** Across the 17 trees reconstructed for the 16 patients, 6 trees were identified as 4-hit events (including B-D), 9 trees were identified as 3-hit events (including the pair of technical replicate from patient ID12565, also see Supplementary Figure S15) and 2 trees were identified as 2-hit events. Samples that contain multiple *k*-th mutants, i.e. multiple possible AML blast genotypes, are marked with a †.

Phylogenies enabled us to reconstruct how AML evolves from initially healthy HSCs. Across the 16 individuals analyzed we found that the AML clones (red stars, Figure 1) most commonly harbour 3 (*n* = 8) or 4 (*n* = 6) driver mutations (Figure 1B-E). The most commonly co-occurring drivers alongside the *DNMT3A* and *NPM1* mutations were in *NRAS* (*n* = 12), *FLT3* (*n* = 11), *TET2* (*n* = 8), *IDH1*/*IDH2* (*n* = 5), *RAD21* (*n* = 4), and *WT1* (*n* = 4) (Supplementary Figure S1-4). In all of the samples where the single-cell level data enabled us to determine the order of mutation acquisition, we found that *DNMT3A* preceded *NPM1. DNMT3A* is typically the first mutation acquired along the evolutionary branch to the AML clone, in agreement with previous findings ^30^. Mutations in the RAS pathway (*FLT3, NRAS*), when observed in our data, are usually late driver events either the 3rd or the 4th “hit”. Acquisition of *NPM1* is usually secondary to *DNMT3A* but earlier than *FLT3* or *NRAS* ^30^. Mutations in *IDH1/2* and *TET2* were found to be mutually exclusive on the ancestral branch of the AML blasts, consistent with previous observations ^31,32^.

The trees also revealed highly diverse patterns of preleukemic evolution. In approximately a third of cases (6/16) we observed evidence of branched evolution indicating multiple evolutionary paths to higher fitness in HSCs. Our data suggests that these parallel competing lineages can coexist for a considerable time as the coalescence events can occur early in a tree. In the remaining 10 samples, we observed patterns of linear evolution which may indicate strong preferences for particular evolutionary paths in the fitness landscape. In addition to the qualitative differences among the trees we also observed clear quantitative diversity at the level of clone sizes. In approximately a third of cases (6/16) we observed that wild-type HSCs – cells carrying no detectable driver mutations – remain the dominant clone in the HSC pool at the time of diagnosis (Figure 1B, C and e.g. ID12565 and ID2261 in E). On the other hand, in 5 cases we observed that the wildtype HSCs have largely been out-competed by later occurring clones carrying multiple driver mutations (Figure 1D, e.g. ID4334 and ID32963 in E). In the remaining samples there appears to be coexistence of intermediate clones – harbouring different numbers of driver mutations – at similar clone sizes (ID47668 and ID26833R in Figure 1E).

### Inferring positive selection from phylogenies

The clone size statistics of the phylogenetic trees are the outcome of clonal competition, mutation and stochasticity during preleukemic evolution. We reasoned that the diverse patterns observed in the AML trees may emerge naturally due to these three interacting elements in the evolutionary dynamics, independent of the specific details of the mutations involved. To test this hypothesis we developed a simplified model of preleukemic evolution which is informed by experimental observations ^8,9,24,33–36^ (Figure 2A, Supplementary note 2).

**Fig. 2.**
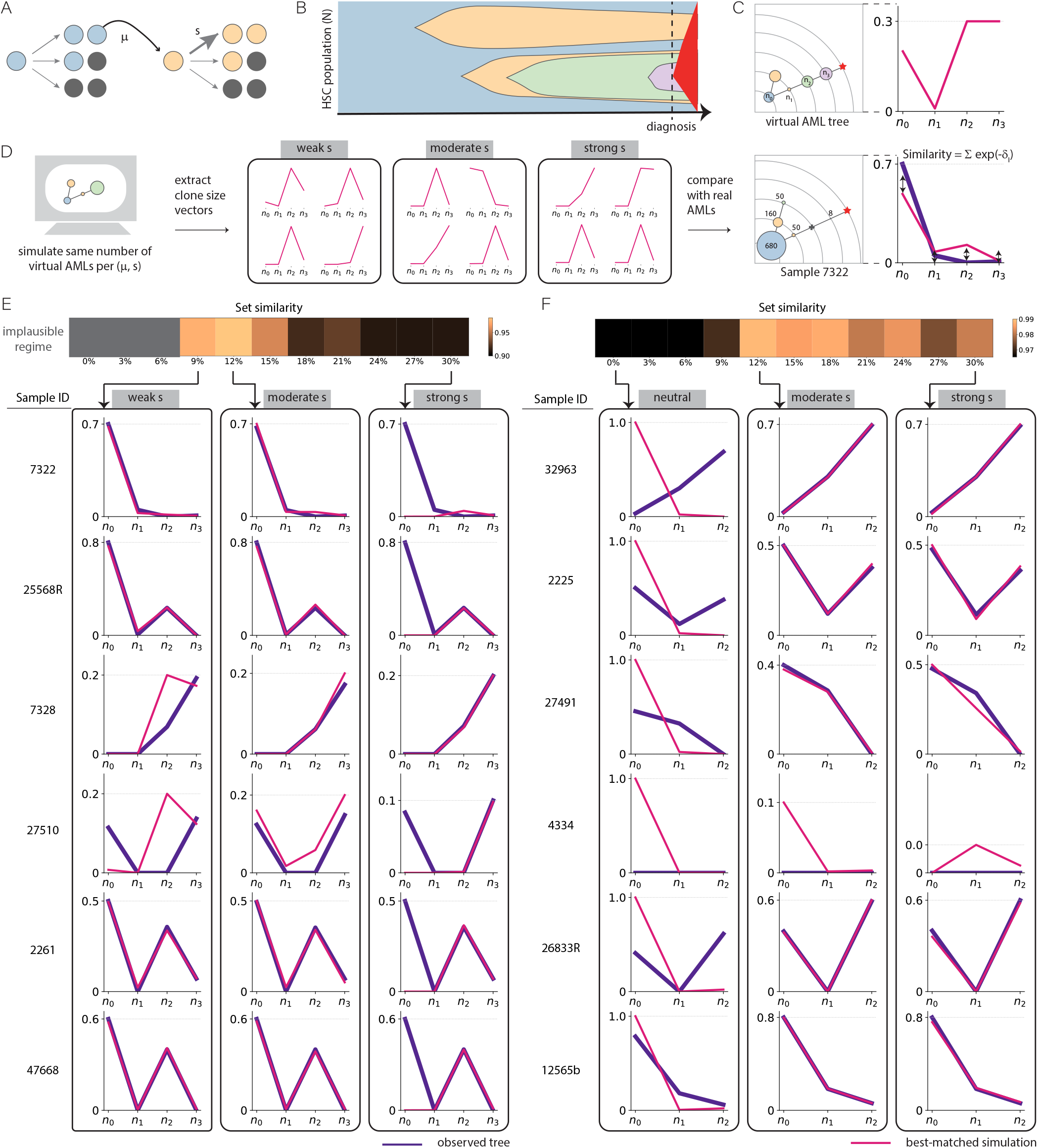
Statistics of preleukemic phylogenetic trees suggests pervasive positive selection operates during preleukemic evolution. **(A)** A model of HSC dynamics as a multi-class branching process where stem cells self-renew symmetrically at rate 1*/τ* and acquire driver mutations stochastically at a rate of *µ* per year. Driver mutations confer a selective advantage *s* per year to the cell. Under the staircase model, every additional mutation confers an additional selective advantage to the lineage. **(B)** An example of the evolutionary dynamics observed in this model where cancer onset is defined by the first lineage that acquires *k* driver mutations with *k* = 4 in this example (red colouring). **(C)** The phylogenetic tree constructed from driver mutations at diagnosis can be used to extract the clone size vector ***n*** =(*n*_0_, *n*_1_, *n*_2_, *n*_3_). **(D)** Stochastic simulations produced four thousand 3-hit and four thousand 4-hit virtual AML events diagnosed before age 80 per parameter combination. The clone size vectors of these simulated ‘virtual AMLs’ (***n***^*′*^, pink lines) were compared against real trees (***n***, purple line) from our single-cell experiments using a similarity metric. **(E, F)** Set similarity (colour scale) between observed trees (purple lines) and closest simulated trees (pink lines) for different selection coefficients across the 12 observed trees considered.

To avoid over-fitting to the data we impose the simplifying assumption that each driver mutation is acquired at the same rate *µ* and confers the same advantage *s* to cells regardless of mutation identity and the effects of driver mutations combine additively. Transformation to AML occurs once *k* driver mutations occur in the same cell (red clone, *k* = 4 case in Figure 2B). This framework can be used to simulate phylogenies of HSCs at the time of transformation. In order to compare the quantitative patterns observed in simulated trees to those experimentally measured in the AML cases, we considered the clone size vector, ***n*** = (*n*_0_, *n*_1_, …, *n*_*k*−1_), which quantifies the clonal frequencies of the wild-type, single mutant, double mutant etc. clones along the main evolutionary branch ancestral to the AML clone at the time of transformation (Figure 2C). The statistical properties of the clone size vector, ***n***, reflect different levels of selection (*s*) operating during preleukemic evolution (Supplementary figure S17).

We used our model to perform in excess of a billion stochastic simulations across a range of fitness effects and mutation rates consistent with previously inferred values ^8,9,13,24,34,37^ (Figure 2D, Supplementary note 2). In these simulations we considered a ‘virtual AML’ to occur when a clone that carries *k* = 3 or *k* = 4 driver mutations emerges before 80 years, in line with the diagnosis ages in our cohort. We recorded 4000 *k* = 3 and 4000 *k* = 4 ‘virtual AMLs’ for every selection strength considered and used them for tree comparison with observed clone size vectors.

Of the sixteen reconstructed trees from our single-cell experiments, twelve were unambiguously genotyped for all driver mutations. We identified the best-matched ‘virtual AML’ for every one of these observed clone size vectors and calculated the total similarity over all the best-matched ‘virtual AMLs’ across a range of selection strengths (Figure 2E and F, Supplementary note 2). This provides an estimate for the likelihood that the set of observed trees is generated from preleukemic evolution driven by the assumed selection strength *s*. We found that the statistical properties of the observed tree set are consistent with simulated tree patterns generated under moderate to strong selection (*s* ≥ 9% per year). These selection strengths cause intermediate clones to expand to high cell fractions and lead to the co-existence of wildtype and multiple mutant clones in the HSC pool at diagnosis. Neutral evolution (*s* = 0%) produces highly homogeneous sets of ‘virtual AML’ tree patterns where the preleukemic stem cell population is always dominated by the wild-type clone. This is inconsistent with the set of our experimentally observed trees, suggesting that stochastic acquisition of mutations alone cannot account for the emergence of AML.

### Validation using an independent single-cell dataset

To test how well these findings generalize to other AML cases, we considered 27 AML trees from an independent singlecell study ^27^. We extracted clone size vectors, ***n***, from each of these cases and performed the same procedure for inferring levels of positive selection (Supplementary note 2). As was observed from our single-cell data, the statistics of the clone size vectors are highly variable with some cases exhibiting the dominance of wild-type clones and others the co-existence of intermediate clones. Consistent with our previous findings, these trees cannot be explained by a model of neutral evolution (left panel, Figure 3). Models of evolution driven by moderate levels of positive selection (9% *< s <* 24% per year) produce clone size vectors that are in close agreement with those observed in the 27 trees (middle panel, Figure 3). Increasing selection strengths even further (*s >* 24% per year) produces clone size vectors which become less consistent with the data (right panel, Figure 3). The selection strengths implied by these data are therefore consistent with the selection strengths inferred from our single-cell data. These findings generalize to cases that were driven by 4 driver mutations (Supplementary Figure S9) as well as trees that show considerable branching (Supplementary Figure S8).

**Fig. 3.**
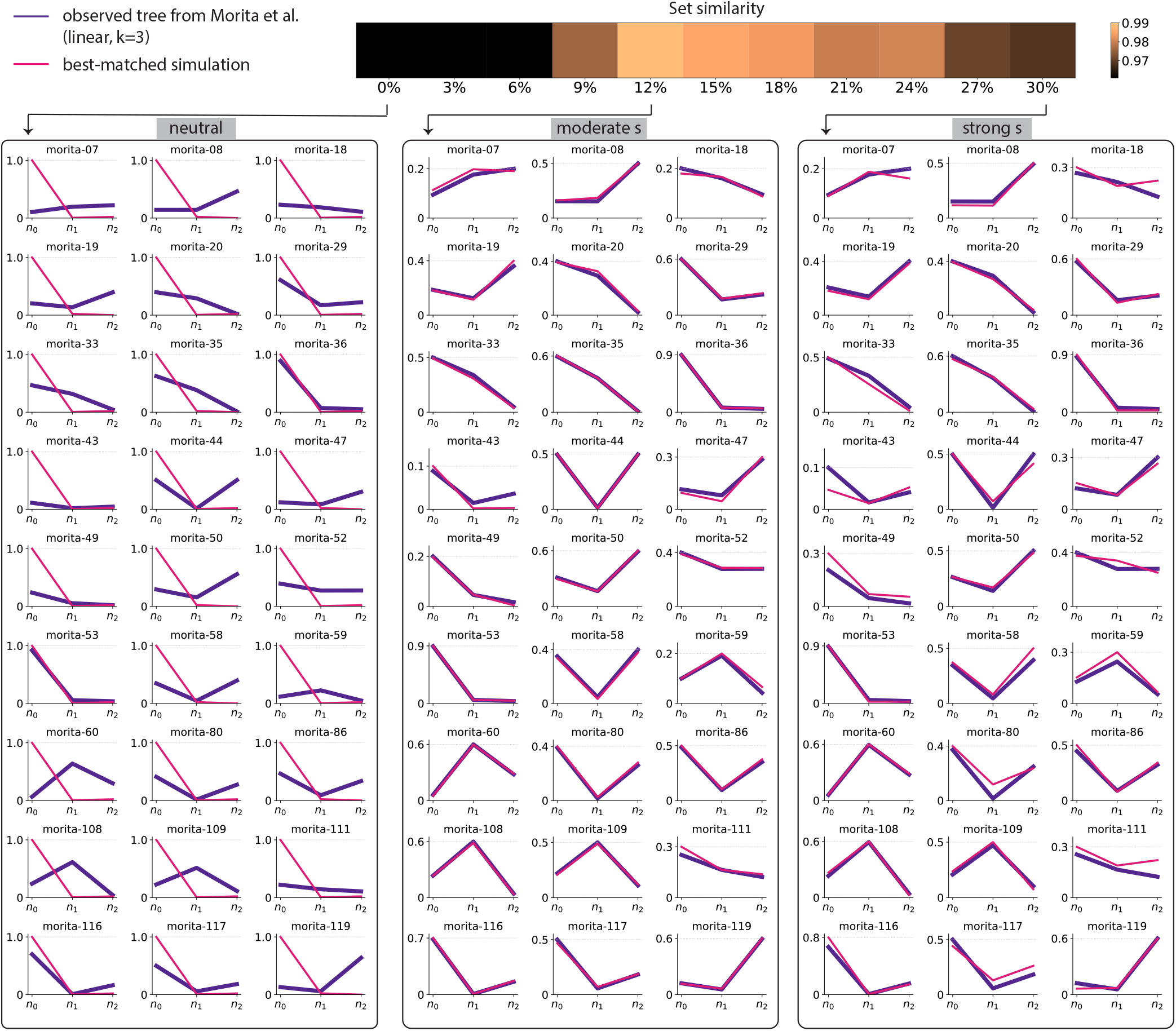
Independent AML single-cell dataset ^27^ validates that moderate selection operates during preleukemic evolution. Clone size vectors from an independent data set of 27 *k* = 3 single-cell AML trees from Morita et al. ^27^ (purple lines) compared to the closest ‘virtual AML’ clone size vectors generated from our stochastic simulations (pink lines) for three different selection strengths (panels). The normalised set similarity (colourscale) between all 27 trees and their closest simulated trees across the full set of selection strengths modelled is shown in the top panel.

### Variation in selection strengths across individuals

To evaluate whether there is evidence of variation in levels of positive selection across individuals, we extracted all *k* = 3 and *k* = 4 AML trees (67 in total) from Morita et al. ^27^ and combined these with our own experimentally observed trees. To estimate the most likely selection levels that drove the preleukemic evolution in each of these 80 trees, we identified the 100 most similar ‘virtual AMLs’ for each case and considered the distribution of the selection strengths that produced them (Figure 4A).

**Fig. 4.**
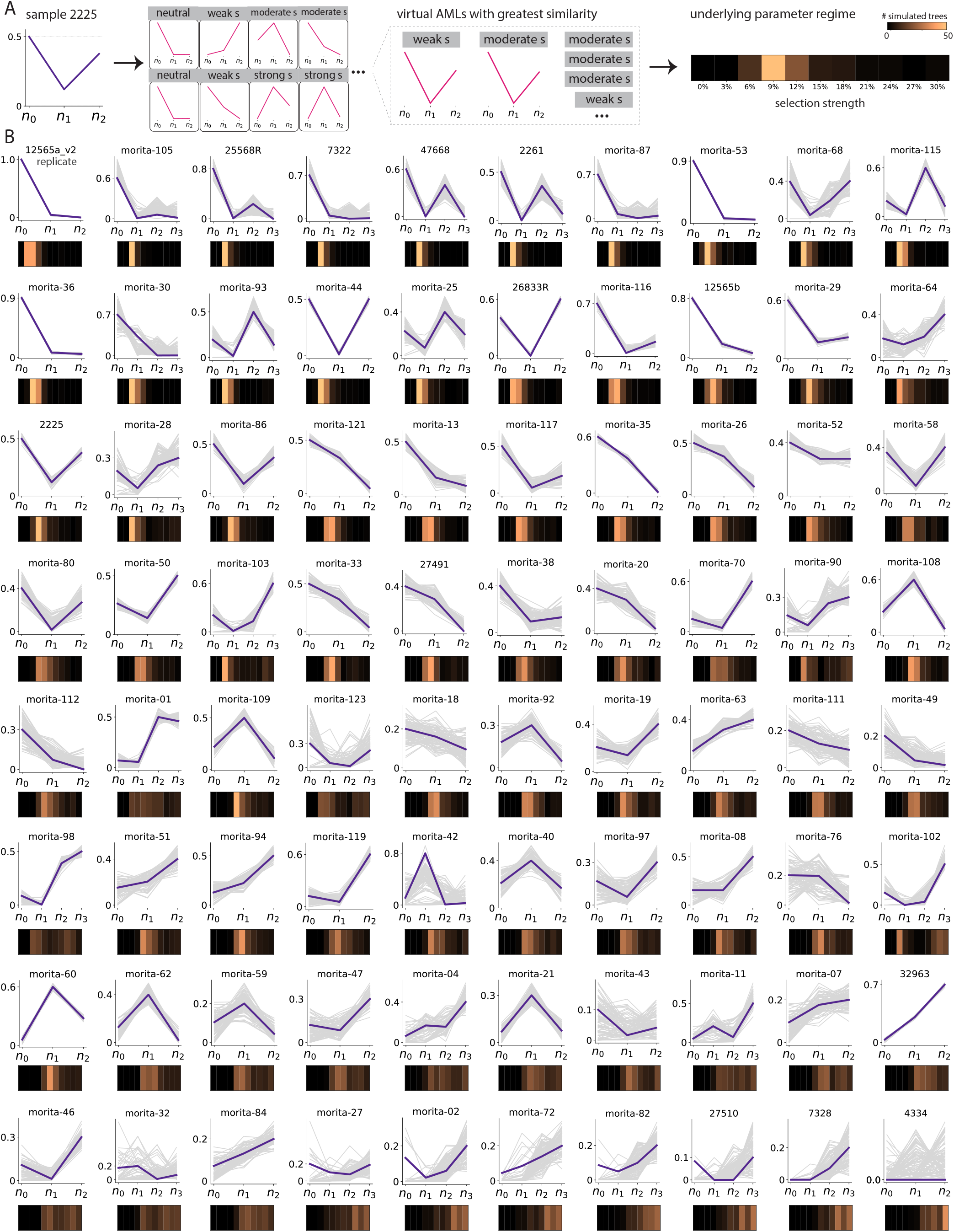
Individual trees show variation in inferred selection levels. **(A)** The 100 ‘virtual AML’ trees highest in similarity were identified for each observed tree. Sample 2225 is shown here. The inferred selection is visualized by the distribution of the number of ‘virtual AMLs’ across the underlying parameter distribution. **(B)** This procedure was carried out for 80 real trees, including 6 four-hit Poon trees, 7 three-hit Poon trees (containing a pair of technical replicate), 44 three-hit Morita trees and 23 four-hit Morita trees.

The majority of trees are consistent with preleukemic evolution driven by selection strengths in the range of *s* = 9% − 24% per year (Figure 4B). However, we observe clear variation in selection strengths across the 80 trees. Trees in which wild-type clones remain dominant at diagnosis (upper rows, Figure 4B) are consistent with weaker selection strengths. In these cases our framework is confident on the inferred levels of selection: the best-matched ‘virtual AMLs’ consistently fall onto a narrow range of ‘*s*’. For example, our inference using this framework strongly suggests that the preleukemic evolution in ID12565 and ID2225 was driven by weak selection strengths at *s* ≈ 3 − 9% per year. Trees with a diverse coexistence of multiple-mutant clones (lower rows, Figure 4B) are inferred to have been driven by stronger levels of positive selection. In these cases, however, the framework is less confident in its inferences: the best-matched ‘virtual AMLs’ come from a wider range of ‘*s*’ (ID32963 and ID7328, Figure 4B).

To assess how finite sampling impacts our inferences, we trained neural networks on subsampled ‘virtual AMLs’ to infer selection strengths *s* and mutation rates *µ* from a single tree (Supplementary note 3). Applying these neural networks to our experimentally observed trees we found that the inferred selection levels are broadly consistent with our previous inferences based on clone-size similarity (Supplementary figure S15). This indicates that subsampling introduced by finite numbers of pHSCs only influences the inferred levels of selection where cell numbers are small.

### In silico modelling identifies features of future AML risk

Our finding that most observed AML trees appear to be consistent with moderate selection strengths suggests that preleukemic clones expand over timescales of decades and could be used to identify individuals at high risk of developing AML. However, a key challenge in identifying robust predictors of risk for rare cancers such as AML is in understanding how rare these features are in individuals not destined to develop the cancer. Using our *in silico* model we are able to compare the evolutionary trajectories that precede ‘virtual AMLs’ (Figure 5A) to the trajectories observed in hundreds of thousands of simulated individuals who never develop ‘virtual AML’. In an HSC population of ∼ 10^5^ stem cells ^8,9^ where driver mutations occur at rate of ∼ 10^−5^ per year ^37^, clones carrying single driver mutations occur routinely in both future AML cases and controls (orange shading, Figure 5B and C). This is consistent with what is known about the prevalence of clonal haematopoiesis in healthy individuals ^14,15^ and suggests that the detection of clones carrying a single driver mutation is unlikely to be a robust predictor of AML. However, the emergence of double mutant clones (i.e. cells harbouring two driver mutations) is predicted to be rare in healthy individuals under the age of 30. In contrast, more than half of future ‘virtual AML’ cases have already acquired a double mutant by the same age (green shading, Figure 5B and C). While the relative risk conferred by the emergence of a double mutant is considerable, its absolute risk remains modest (Figure 5D) (see Supplementary Figure S16 for *k* = 3). The emergence of a triple mutant clones before the age of 50 confers both a large relative and absolute risk of developing a four-hit ‘virtual AML’ before age 70.

**Fig. 5.**
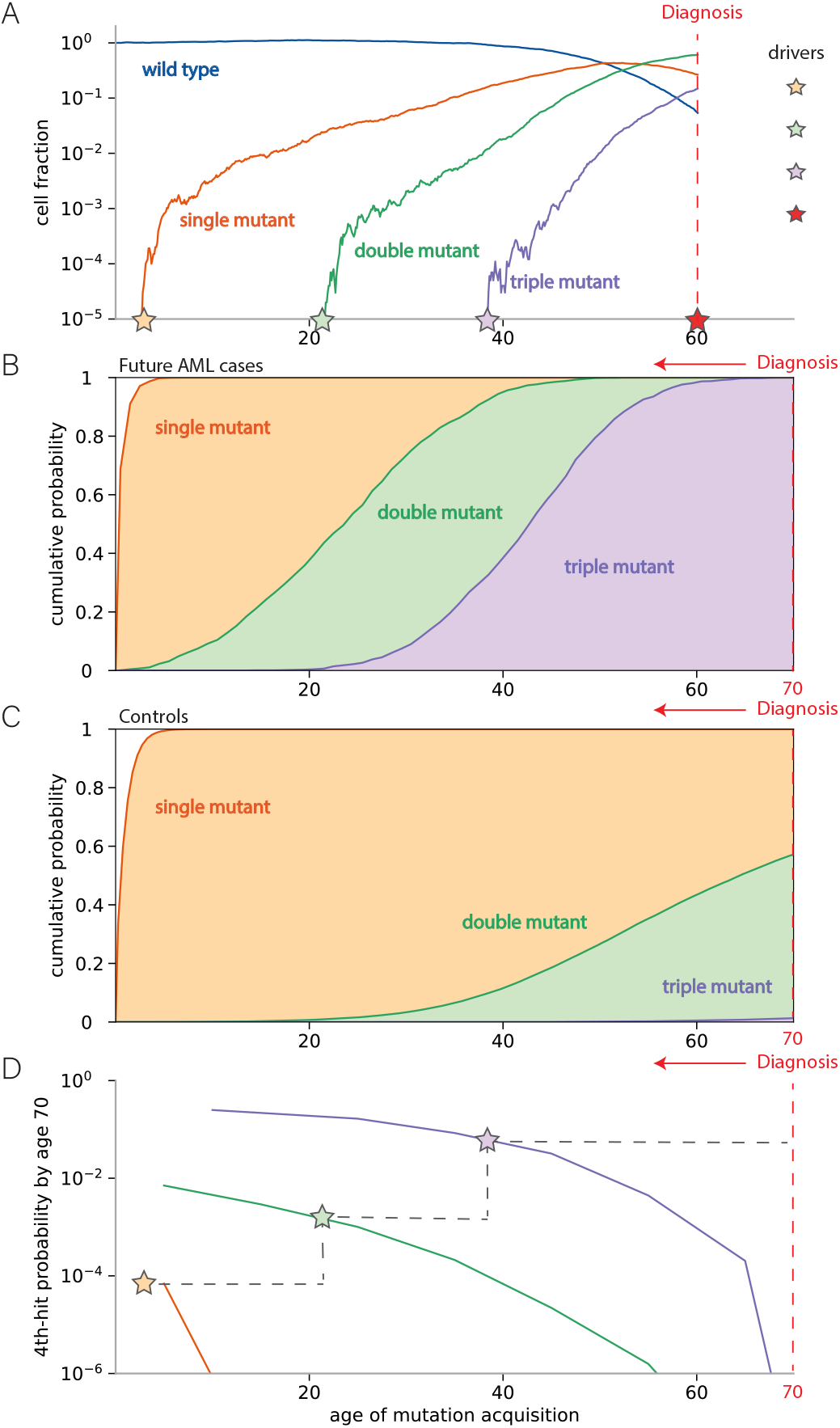
Early double-mutant and triple-mutant clones identify a highrisk group for AML transformation. **(A)** An example of the preleukemic clonal history of a simulated individual who acquires an AML-defining 4th-hit at age 60 where AML drivers confer moderate levels of positive selection. **(B)** Distribution of mutation acquisition times in simulated individuals (under moderate selection and at medium mutation level) who acquired 4th-hit before age 70. **(C)** Distribution of mutation acquisition times in unselected simulated individuals, showing marked differences compared to simulated individuals destined to develop AML. **(D)** The absolute risk of AML transformation before age 70 is dependent on when single (orange), double (green) and triple (purple) mutant clones are acquired. This is illustrated for the individual in (A) for whom the absolute risk jumps upwards (grey dashed line) whenever the individual acquired a further mutation (star).

In summary, by simulating the evolutionary dynamics using parameters inferred from the experimental trees and examining large numbers of virtual AML cases and controls one can learn the general features which identify future AML cases. Multiple-mutant clones (carrying >1 driver mutation in the same lineage) that arise anomalously early (e.g. in individuals <30 years of age) carry a considerable risk of transforming to AML.

## Discussion

We have shown that preleukemic clones, which harbour subsets of the driver mutations present in the AML clone, coexist in the HSC/multipotent progenitor (MPP) pool at the time of AML diagnosis. Single-cell sequencing of these clones enabled the construction of phylogenetic trees depicting the clonal evolution that occurs in HSCs antecedent to AML. We found that clone size patterns revealed by the phylogenetic trees were highly variable across individuals. Using evolutionary modelling we showed that these diverse patterns are representative outcomes of somatic evolution that is driven by pervasive positive selection. The major implication of these inferred selection strengths is that preleukemic clones expand from low frequencies to become detectable decades prior to the AML transformation event. This provides a long window of opportunity for risk prediction and potential intervention. Our inferences are based on the assumption that selection strengths remain constant throughout life. However we cannot rule out that selection pressures may vary in time due to extrinsic factors including ageing ^34,38^, cytotoxic therapies and smoking ^37^.

There are a number of technical limitations in our work that need to be considered when interpreting the results. Chief amongst these is our premise that the sizes of preleukemic clones are determined only by the forces of selection, mutation and drift operating during preleukemic evolution. One important possible technical effect is that clones sizes may also reflect biases introduced during the experimental process of isolating and sequencing HSCs. Specifically, if the immunophenotype of the HSC co-varies consistently with the presence of a particular driver mutation, this could cause systematic biases in our inference of selection strengths. To evaluate the influence of this effect we performed a technical replicate of our experimental procedure on the bone marrow aspirates from one patient (ID12565 in Supplementary Figure S15) and confirmed that our model inferences were unaffected. The validation of our inferences on an independent single-cell dataset ^27^ generated from unsorted samples also suggests that the experimentally observed clone size vectors are not primarily dominated by systematic biases introduced during the isolation of pHSCs.

Whilst the conceptual model presented here is clearly an oversimplification of the underlying biology (neglecting epistasis between driver mutations, extrinsic perturbations altering the fitness of clones ^34,39^, explicit effects of ageing ^38^ and many other factors) it nevertheless highlights that the patterns observed in the phylogenetic trees can be explained by a far simpler model which has as its defining feature the requirement for positive selection on intermediate clones. In order to focus on the general behaviours that emerge from positive selection, the model intentionally ignores differences between driver mutations (i.e. all drivers confer the same selective advantage to HSCs and their effects combine additively in fitness effect). This simplified model highlights that the divergence between healthy and pre-malignant evolutionary dynamics in blood usually occurs due to the emergence of an anomalously early double mutant clone. We propose that the detection of rare clones harbouring two driver mutations within a single cell in young adults could provide a rational basis for identifying individuals at high risk of progression to AML. This conceptual framework may also provide a potential route to resolving the apparent discrepancies between models of oncogenesis based on stochastic mutation accumulation and bad luck ^6,40–42^, the detection of driver mutations in healthy tissues ^8,9,14–17,43–45^ and the observed lifetime risk of AML (Supplementary Figure S18).

## Acknowledgements

We thank Daniel Fisher for helpful discussions and the team at Mission Bio for their technical support throughout this project. We thank all members of the Blundell lab for input. Y.P.G.P and J.R.B are funded by a UKRI Future Leaders Fellowship and by the CRUK Cambridge Cancer Centre. We thank Cambridge NIHR BRC Cell Phenotyping Hub for flow cytometry sorting. E.L. is funded by a Wellcome – Royal Society Sir Henry Dale Fellowship 107630/Z/15/Z and A.V. by Gates Cambridge Trust PhD Scholarship (10350885) until 2022. Work in E.L. group is funded by BIRAX (47BX16ELLS) and core support grants by Wellcome and Medical Research Council (MRC) to the Wellcome-MRC Cambridge Stem Cell Institute 203151/Z/16/Z and the UKRI Medical Research Council (MC_PC_17230). For the purpose of open access, the author has applied a CC BY public copyright licence to any Author Accepted Manuscript version arising from this submission.

## Data and code availability

All code used in this study are available on the [Blundell lab Github page](https://github.com/blundelllab). The raw sequencing data will be deposited on the European Nucleotide Archive.

## Supplementary Note 1: Experimental Methods

### Sample selection

Bone marrow aspirate samples containing mononuclear cells were collected between 1998-2017 from treatment-naive *de novo* AML patients either enrolled in the Dutch-Belgian Cooperative Trial Group for Hematology-Oncology (HOVON) or the Swiss Group for Clinical Cancer Research (SAKK) clinical trials. All of them had provided written informed consent in accordance with the Declaration of Helsinki. We processed 17 samples from 16 patients harbouring both *DNMT3A* and *NPM1* hotspot mutations, two of which being biological replicates (ID12565a and 12565b). Patients’ ages range between 37 − 75 at the time of sample collection (labelled on Supplementary Figure S1, S2, S3, S4).

### AML blast genotyping and phenotyping

High molecular weight genomic DNA was isolated from bone marrow samples or peripheral blood. Bulk gene re-sequencing was performed on the diagnostic samples using a 54 gene panel containing the most frequently mutated genes in myeloid malignancies. Targeted next-generation sequencing (NGS) was carried out with the Illumina TruSight Myeloid Sequencing panel (HOVON-SAKK). The genomic libraries were sequenced on the Illumina platform (Illumina, San Diego, CA) with mean target coverage of at least 500x and an average sequencing depth of 2500x. Variant calling was carried out as described previously ^46^. This procedure enables the identification of high-confidence driver mutations during preleukemic evolution. Variants called that were between 20 − 80% VAF (80 − 100% VAF if homozygous) and considered highly pathogenic were assigned to the high-confidence set of driver mutations for that sample.

### Isolation of pHSCs

BM mononuclear cells samples listed in Supplementary Table 1 were thawed using 50% IMDM/50% FCS with 1:100 DNase and MNCs were counted manually using a haemacytometer. Positive selection of CD34+ cells from the remaining sample was achieved by incubating with CD34 Micro Beads (30µL/10^8^ cells, Miltenyi), FcR Blocking Reagent (30µL/10^8^ cells) and PBS + 3% FBS (90µL/10^8^ cells) for 30 minutes at 4^*°*^C. Cells were then washed and resuspended in MACS buffer and applied to a prepared LS magnetic column for manual selection as per the Miltenyi user manual. Samples with 1-5×10^7^ MNCs were selected using EasySep (StemCell Technologies). EasySep was performed by incubating in the selection cocktail for 15min and with magnetic particles for 10min at room temperature. Once in the magnet, supernatant containing CD34-cells was poured out and frozen separately, while CD34+ cells were separated for staining and selection.

**Table 1.**
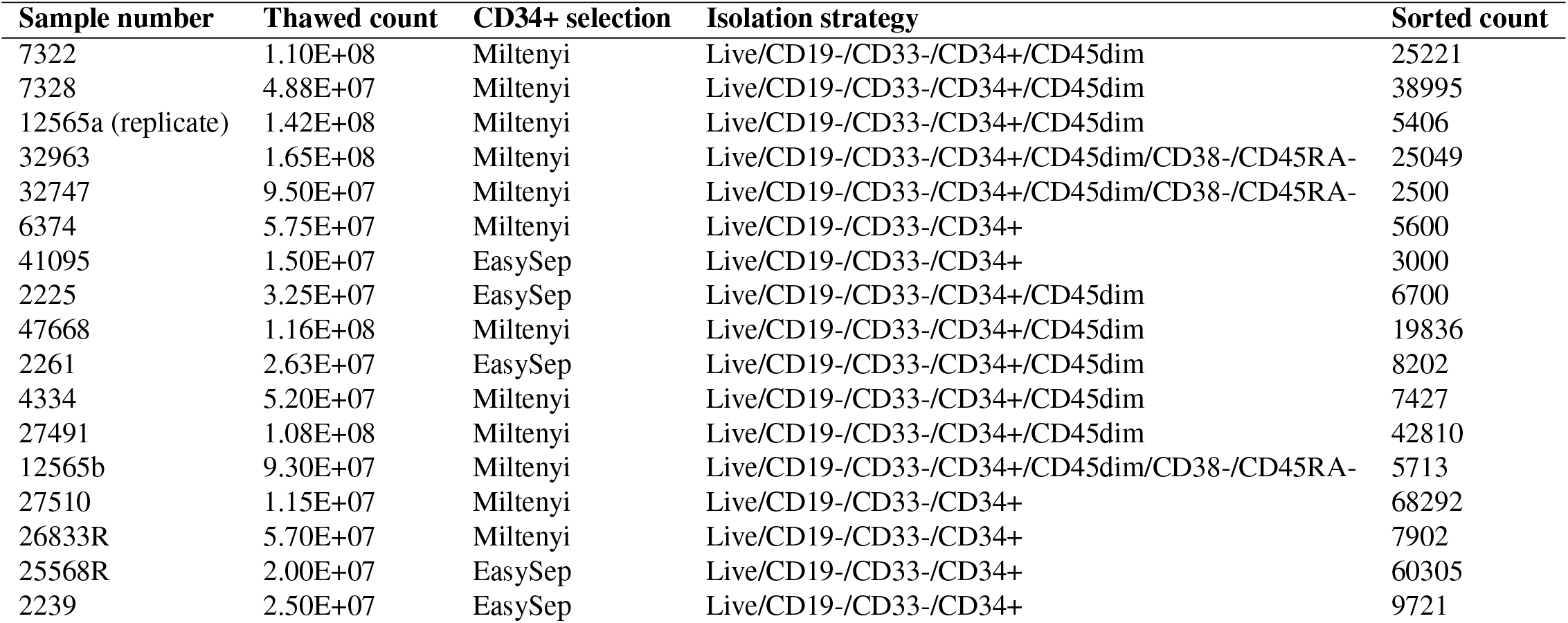
PHSC isolation strategies.

MNCs were then stained for flow cytometry using the antibody panel in Table 2. We then performed fluorescence-activated cell sorting (FACS) on the CD34+ cells in order to isolate preleukemic HSCs (pHSCs) in bulk. Here we define pHSCs based on being phenotypically-normal HSCs in AML samples. These pHSCs should only harbour early driver mutations, a subset of the leukemic-specific mutations. We aimed to isolate the CD19−/CD33−/CD34+/CD38−/CD45dim/CD45RA-compartment and gating requirements were relaxed in some cases to meet cell input requirements for the single-cell sequencing platform (Supplementary Figure S5). All pHSC samples were at a minimum, CD19−/CD33−/CD34+, and some were also CD45dim. Any leukemic stem cells (red stars in relevant figures) leaked into the sorted population could be identified based on cell genotype and were excluded in downstream analysis. The sorted population size varies between 2500 − 68292 cells across the 17 samples as a result of variable sample immunophenotype on blasts, blast percentage and viable cell count after thawing.

**Table 2.** List of all antibodies used for FACS in this study.

| Antibody | Dilution | Manufacturer | Catalog # |
| --- | --- | --- | --- |
| Mouse monoclonal anti-CD45RA, Alexa 700 | 1:300 | BioLegend | 304120 |
| Mouse monoclonal anti-CD33, APC | 1:200 | BD Biosciences | 345800 |
| Mouse monoclonal anti-CD34, APC-Cy7 | 1:100 | BioLegend | 343514 |
| Mouse monoclonal anti-Zombie, Aqua (BV510) | 1:800 | BioLegend | 423101 |
| Mouse monoclonal anti-CD45, BV785 | 1:100 | BioLegend | 304048 |
| Mouse monoclonal anti-CD19, FITC | 1:100 | BioLegend | 302206 |
| Mouse monoclonal anti-CD90, PE | 1:50 | BioLegend | 328110 |
| Mouse monoclonal anti-CD49f, PE-Cy5 | 1:100 | BD Biosciences | 551129 |
| Mouse monoclonal anti-CD38, PE-Cy7 | 1:100 | BioLegend | 303516 |

The sorted pHSC population was then put through the commercial microfluidic-based encapsulation platform (‘Tapestri’, by Mission Bio) during which genomic DNA was barcoded and amplified within single-cell droplets. The panel used was the catalogue myeloid panel which consists of 312 amplicons (65 kb) that target 45 commonly mutated myeloid genes ^47^. Samples were then made into libraries and over-sequenced using standard paired-end (2 *×* 150 bp) sequencing on Illumina platforms (MiSeq and NovaSeq 6000 with the SP flowcell). Sequencing read output shows that > 5% single cells were captured per sample but typically only ≈ 10 − 1000 single cells were well genotyped at all driver mutation positions (Supplementary Figure S1, S2, S3, S4).

### Single-cell variant calling

FASTQ files generated were then uploaded to the Tapestri DNA pipeline v2 after downsizing for a target coverage of 50x per amplicon per cell. Adapter sequences were trimmed from sequenced reads using Cutadapt v2.3^48,49^ and reads were mapped to the hg19 reference genome using the BWA-MEM algorithm ^50,51^. Barcodes within a small Hamming or Levenshtein distance from corrupted barcodes were corrected. Genome Analysis Toolkit ^52^ with a joint calling approach that follows GATK Best Practices recommendations ^53,54^ was used to call single nucleotide variants and indels. A custom genotyping method was used to target internal tandem duplications in the *FLT3* gene in the Tapestri pipeline. However, *FLT3-ITD* was not detected in our samples due to their low variant allele frequencies even when present (ID 32747). A final .loom file was generated for each sample, which contains a matrix of values detailing the allele frequencies of variants called. The data was then visualized using the commercial software Tapestri Insight v2.2 provided by Mission Bio and analyzed quantitatively in Python 3.10.9.

### Dropout clone analysis and missing genotypes

The dropout rates estimated based on known heterozygous variants ranges between 6 − 26% (average≈ 14%) across 15 samples (estimates not available for the remaining two). This indicates that on average 14% of cells genotyped for a certain variant was wrongly labelled as either harbouring a homozygous variant or the reference allele. Due to this effect, not all clonal genotypes detected are true and some may be dropout clones from true clones. These clones were marked clearly on the phylogenetic trees and only the total cell numbers combined across ambiguous clones were reported (ID32747, ID41095, ID12565a and ID2239) when the complete mutation order cannot be ascertained. In cases where the mutation orders were clear from the bulk mutation VAFs we resolved dropout clones by assigning cells of dropout clones back to their true clones noting that dropout clones are always considerably smaller than the true clones they belong to.

### Construction of phylogenies

Clone sizes measured from single-cell DNA sequencing were used to construct phylogenetic trees for each sample. Cells were assigned to respective levels based on the number of high-confidence driver mutations they harbour. There are two reasons any high-confidence drivers may not be recovered during single-cell DNA sequencing. First it is possible that the panel does not cover the driver mutation or that the variant position was not genotyped. Bulk VAFs of these high-confidence driver mutations are marked with red triangle markers. They are not included in the phylogenetic tree unless they are *DNMT3A* or *NPM1* mutations. Samples harbouring high-confidence driver mutations whose presence cannot be ascertained are labelled with ‘hidden driver’. Second the high-confidence driver mutation may also be genuinely missing in the preleukemic HSC pool, in which case the cell fraction of the driver mutation (purple bars) is shown as zero. The assumption is that these mutations caused the AML transformation event and belong to the blast genotype (e.g. in sample ID12565a, ID2225, 32747, 7328, 7322). Clone size vectors were then extracted from the reconstructed phylogenetic trees where only the clonal sizes on the ancestral branch of the AML and of side mutants up to k-th hit (not including blast cells) were considered.

#### Sample 7328

Patient age was 68 years (Supplementary figure S1A). (DNMT3A R882H -> NPM1 W288Cfs* -> NRAS G12S -> IDH2 R140Q) formed the set of high-confidence drivers for this patient, according to the bulk sequencing results. This sequence of mutation acquisition is consistent with observed clonal genotypes in the single cell sequencing results. We assigned cells to their respective clones based on mutation co-occurrence observed.

#### Sample 7322

Patient age was 63 years (Supplementary figure S1B). (DNMT3A R882H, NPM1 W288Cfs*, CBL C419F, ETV6 R55fs, TET2 A295fs, RAD21 splicing) formed the set of high-confidence drivers for this patient, according to the bulk sequencing results. TET295fs was not genotyped in the single cell sequencing due to poor amplicon performance and RAD21 splicing was not covered by the panel (hence sample marked with ‘hidden driver’). The acquisition order of the remaining four high-confidence drivers was determined by observed clonal genotypes in the single cell sequencing results. We also identified an additional branch of evolution in the preleukemic stem cell pool ASXL1 Q829* -> TET2 C1271Lfs*. Both mutations are at much lower variant allele frequencies in the DNA sequencing results of the bulk sample.

#### Sample 32963

Patient age was 62 years (Supplementary figure S1C). (DNMT3A W753C, NPM1 W288Cfs*, TET2 L371X, TET2 H724fs) formed the set of high-confidence drivers for this patient, according to the bulk sequencing results. TET2 H724fs was not covered by the panel (hence sample marked with ‘hidden driver’). TET2 L371X was genotyped at very low cell fraction in the preleukemic stem pool due to sorting against blast phenotype, consistent with the mutation acquisition order DNMT3A W753C -> NPM1 W288Cfs*-> TET2 L371X.

#### Sample 6374

Patient age was 55 years (Supplementary figure S1D). (DNMT3A R771X -> NPM1 W288fs*) formed the set of high-confidence drivers for this patient, according to the bulk sequencing results. This sequence of mutation acquisition is consistent with observed clonal genotypes in the single cell sequencing results.

#### Sample 2225

Patient age was 68 years (Supplementary figure S2A). (DNMT3A R882H, NPM1 W288Cfs*, IDH1 R132H) formed the set of high-confidence drivers for this patient, according to the bulk sequencing results. The acquisition order of the high-confidence drivers was determined by observed clonal genotypes in the single cell sequencing results.

#### Sample 32747

Patient age was 55 years (Supplementary figure S2B). (DNMT3A R882H, NPM1 W288Cfs*, IDH2 R140Q) formed the set of high-confidence drivers for this patient, according to the bulk sequencing results. NPM1 W288fs* was not genotyped in the single cell sequencing results due to poor amplicon performance (hence sample marked with ‘hidden driver’). IDH2 R140Q was not detected, suggesting that it belongs to the blast genotype. Wild type cells were assigned under the assumption that DNMT3A R882H was the first mutation acquired.

#### Sample 41095

Patient age was 56 years (Supplementary figure S2C). (DNMT3A E426X, NPM1 W288Cfs*, RAD21 N190fs) formed the set of high-confidence drivers for this patient, according to the bulk sequencing results. DNMT3A E426X was not covered by the panel and RAD21 N190fs was not genotyped in single-cell sequencing. Order of mutation acquisition cannot be ascertained.

#### Sample 12565a and b

Patient age was 50 years (Supplementary figure S2D and S4C). Two technical replicates had been processed for this sample. (DNMT3A R882H, NPM1 W288Cfs*, IDH2 R140Q) formed the set of high-confidence drivers for this patient, according to the bulk sequencing results. This sequence of mutation acquisition was clear from observed clonal genotypes in the single cell sequencing results of sample 12565b. We assigned cells to their respective clones based on mutation co-occurrence observed. NPM1 W288Cfs* was not genotyped for sample 12565a and therefore cells belonging to the single and double mutant clones cannot be assigned with certainty.

#### Sample 25568R

Patient age was 75 years (Supplementary figure S3A). (DNMT3A G706E, NPM1 W288Cfs*, NRAS G12D, TET2 S657X, RAD21 Q293fs) formed the set of high-confidence drivers for this patient, according to the bulk sequencing results. All drivers were well-genotyped in single-cell sequencing except for RAD21 Q293fs (hence sample marked with ‘hidden driver’). We assigned cells to their respective clones based on mutation co-occurrence observed.

#### Sample 27491

Patient age was 59 years (Supplementary figure S3B). (DNMT3A R882C, NPM1 W288Cfs*, PTPN11 E76G, PTPN11 T73I) formed the set of high-confidence drivers for this patient, according to the bulk sequencing results. The sequence of mutation acquisition is clear from observed clonal genotypes in single cell sequencing results. We have assumed that DNMT3A and NPM1 mutations alone are insufficient for blast transformation.

#### Sample 2261

Patient age was 37 years (Supplementary figure S3C). (DNMT3A R882H, NPM1 W288Cfs*, FLT3 D835Y, SMC3 R661P) formed the set of high-confidence drivers for this patient, according to the bulk sequencing results. This sequence of mutation acquisition is clear from observed clonal genotypes in the single cell sequencing results. We assigned cells to their respective clones based on mutation co-occurrence observed.

#### Sample 4334

Patient age was 54 years (Supplementary figure S3D). (DNMT3A delins, NPM1 W288Cfs*, NRAS G12V) formed the set of high-confidence drivers for this patient, according to the bulk sequencing results. We only found one single clone in the single cell sequencing results. It is not ancestral to the blasts and diverged from the main branch earlier on. The clone size vector used for downstream analysis is (0,0,0).

#### Sample 2239

Patient age was 48 years (Supplementary figure S3E). (DNMT3A M548I, NPM1 W288Cfs*, FLT3 D835Y) formed the set of high-confidence drivers for this patient, according to the bulk sequencing results. DNMT3A M548I was not genotyped in our single cell sequencing. Assuming DNMT3A mutation was the first driver, we assigned cells to double mutants.

#### Sample 27510

Patient age was 56 years (Supplementary figure S3F). (DNMT3A R736H, NPM1 W288Cfs*, SMC3 R381Q, FLT3 D835V) formed the set of high-confidence drivers for this patient, according to the bulk sequencing results. The sequence of mutation acquisition can be inferred from observed clonal genotypes (with some ambiguity, as labelled) in single cell sequencing results.

#### Sample 47668

Patient age was 71 years (Supplementary figure S4A). (DNMT3A P115S, NPM1 W288Cfs*, TP53 ins, TET2 V781fs, TET2 Q1539X, SRSF2 P95L) formed the set of high-confidence drivers for this patient, according to the bulk sequencing results. SRSF2 P95L and TET2 Q1539X were not genotyped in the single cell sequencing (hence sample marked with ‘hidden driver’). The acquisition order of the remaining four high-confidence drivers was inferred (with some ambiguity, as labelled) from observed clonal genotypes.

#### Sample 25568R

Patient age was 70 years (Supplementary figure S4B). (DNMT3A L584H, NPM1 W288Cfs*, WT1 R363fs, RAD21 A572fs) formed the set of high-confidence drivers for this patient, according to the bulk sequencing results. All drivers were well-genotyped in single cell sequencing except for RAD21 A572fs (hence sample marked with ‘hidden driver’). We assigned cells to their respective clones based on mutation co-occurrence observed.

**Fig. S1.**
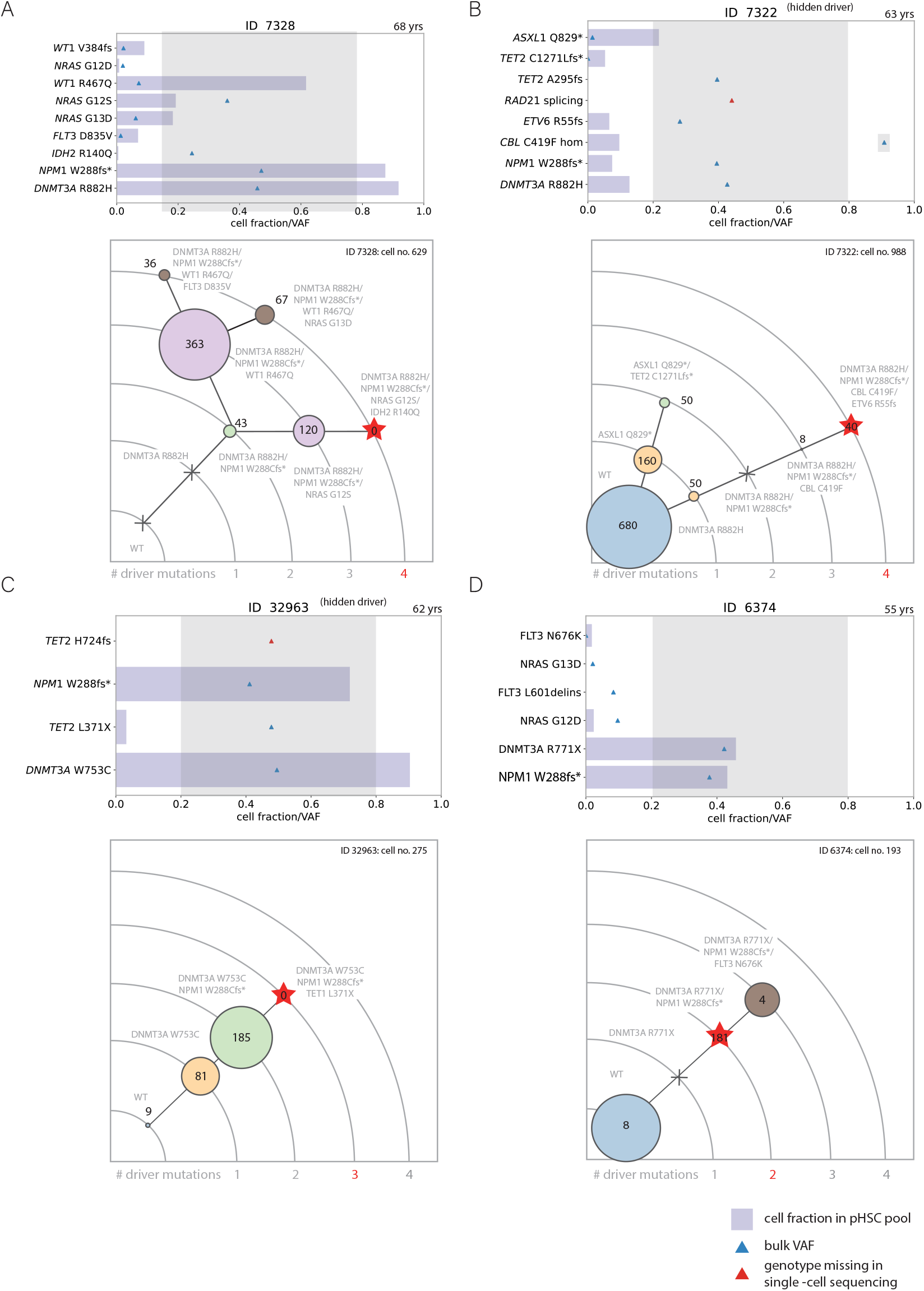
Single-cell DNA sequencing was used to construct pHSC phylogenetic trees for individual patients (first batch - 1). **(A-D)** Variant information from bulk sample (triangle) and cell fractions detected in single cell sequencing (purple horizontal bar) were used to construct phylogenetic trees for the preleukemic HSC population in each patient. Variants falling within VAF 0.2 − 0.8 (VAF≥ 0.9 for homozygous) in bulk were considered as part of the high-confidence driver mutation set and the number of these drivers harboured by a clone was used to assign the clone to corresponding arc levels in the phylogenetic tree. If variants were not covered by the single-cell sequencing panel or not genotyped, the order of acquisition for those mutations cannot be ascertained. These samples are labelled with ‘hidden driver’. Clonal genotypes and sizes were labelled on the constructed phylogenetic trees of each sample. Sizes of circles are proportional to clonal sizes and extinct clones are indicated with grey crosses.

**Fig. S2.**
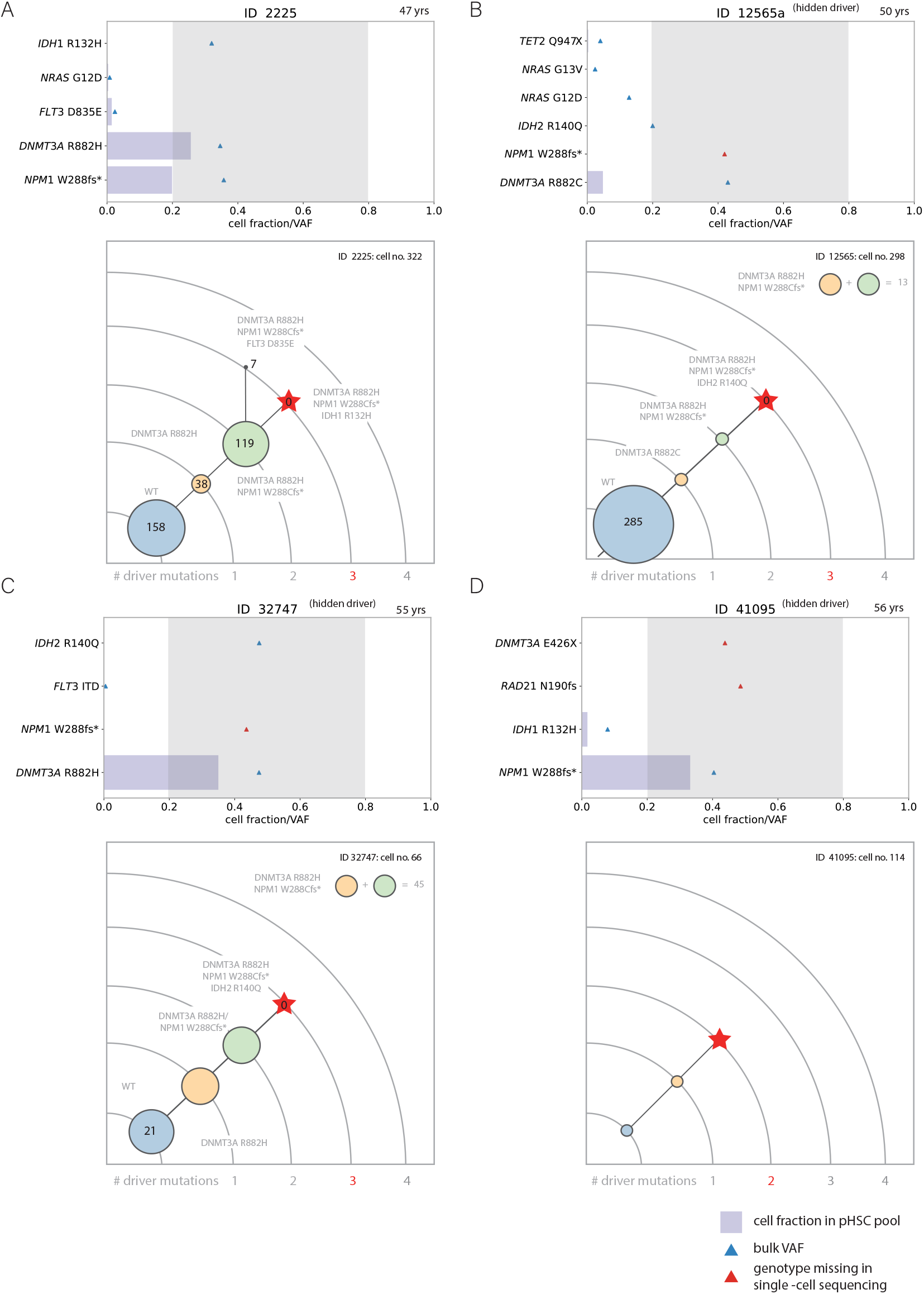
Single-cell DNA sequencing was used to construct pHSC phylogenetic trees for individual patients (first batch - 2). **(A-D)** Variant information from bulk sample (triangle) and cell fractions detected in single cell sequencing (purple horizontal bar) were used to construct phylogenetic trees for the preleukemic HSC population in each patient. Variants falling within VAF 0.2 − 0.8 (VAF≥ 0.9 for homozygous) in bulk were considered as part of the high-confidence driver mutation set and the number of these drivers harboured by a clone was used to assign the clone to corresponding arc levels in the phylogenetic tree. If variants were not covered by the single-cell sequencing panel or not genotyped, the order of acquisition for those mutations cannot be ascertained. These samples are labelled with ‘hidden driver’. Only the total clonal sizes summed across the ambiguous clones were reported (upper right corner equations in tree plots). Clonal genotypes and sizes were labelled on the constructed phylogenetic trees of each sample. Sizes of circles are proportional to clonal sizes and extinct clones are indicated with black crosses.

**Fig. S3.**
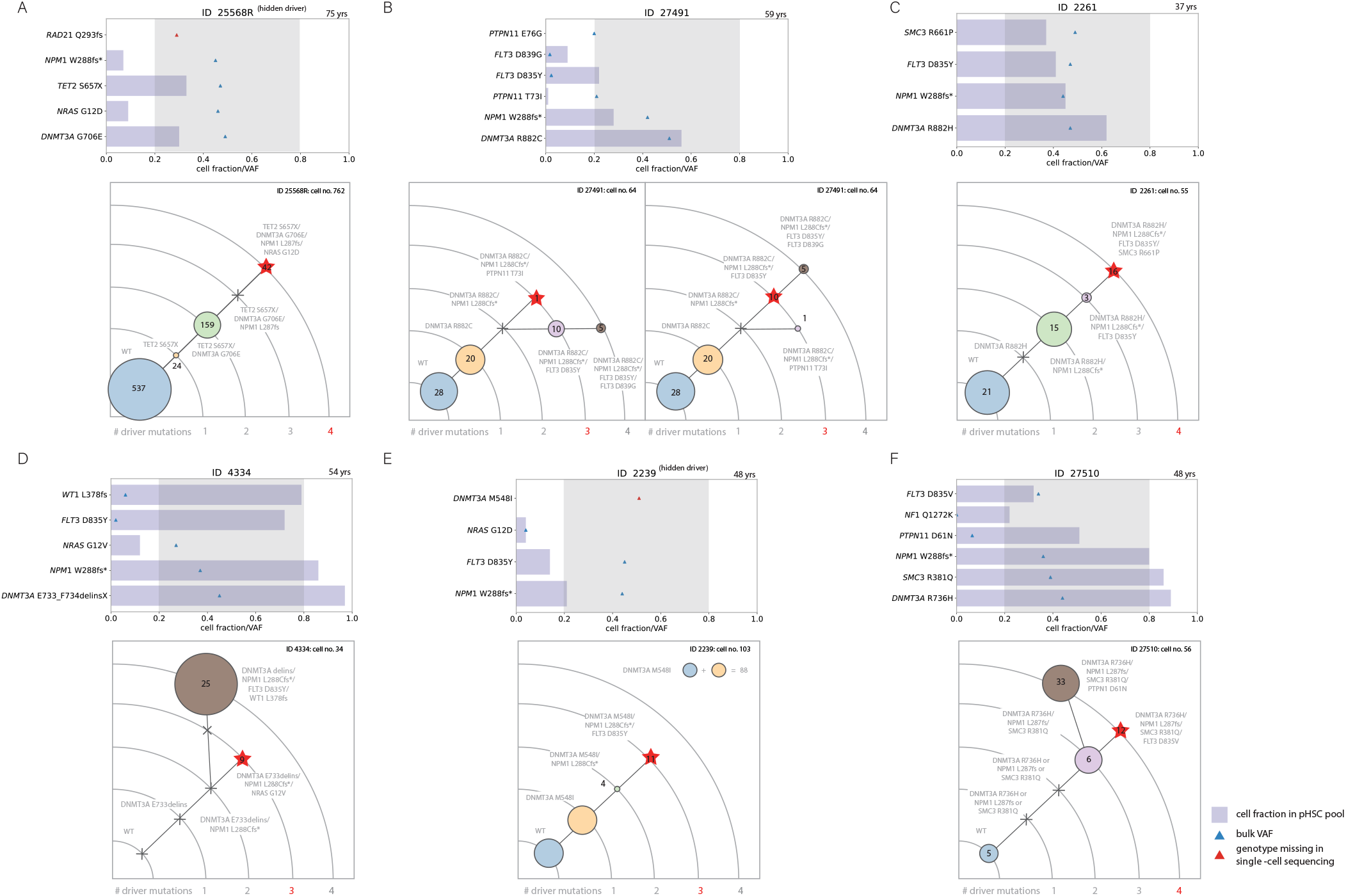
Single-cell DNA sequencing was used to construct pHSC phylogenetic trees for individual patients (second batch). **(A-F)** Variant information and constructed phylogenies are shown for six additional samples. For ID27491 there are two possible phylogenetic trees both of which produce similar clone size vectors.

**Fig. S4.**
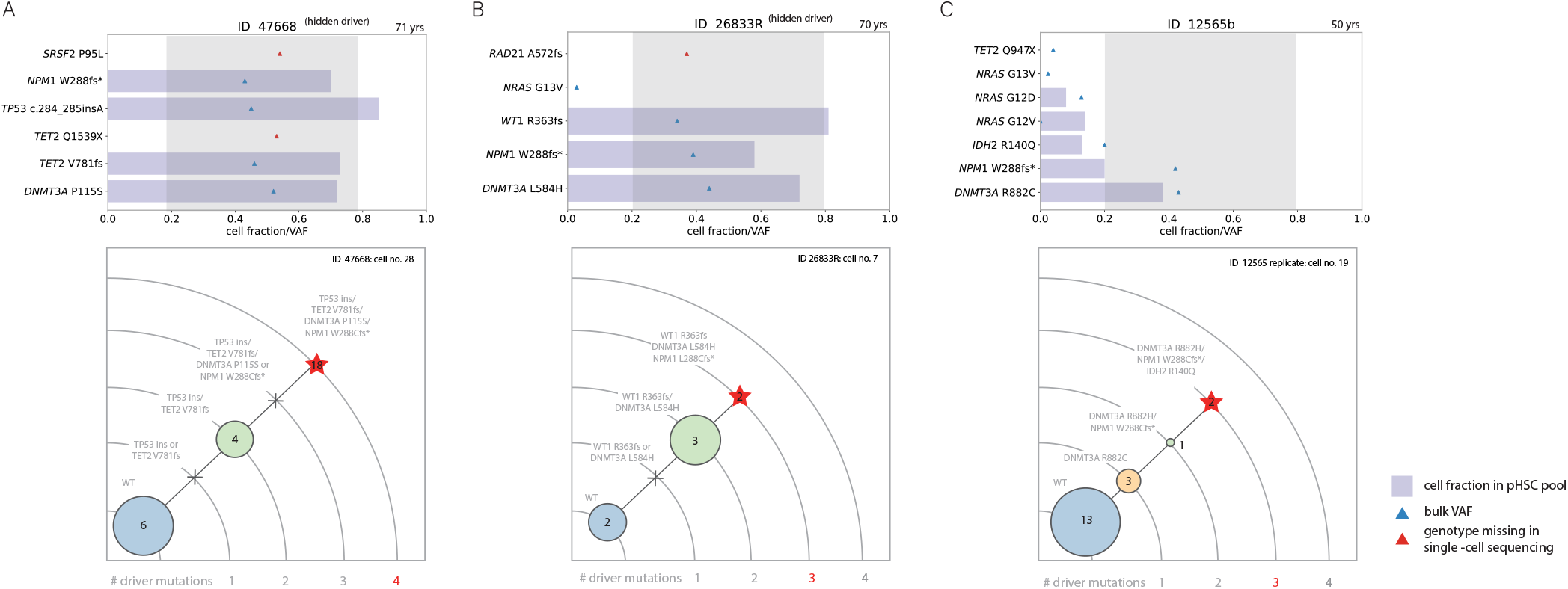
Single-cell DNA sequencing was used to construct pHSC phylogenetic trees for individual patients (second batch). **(A-C)** Variant information and constructed phylogenies are shown for three additional samples.

**Fig. S5.**
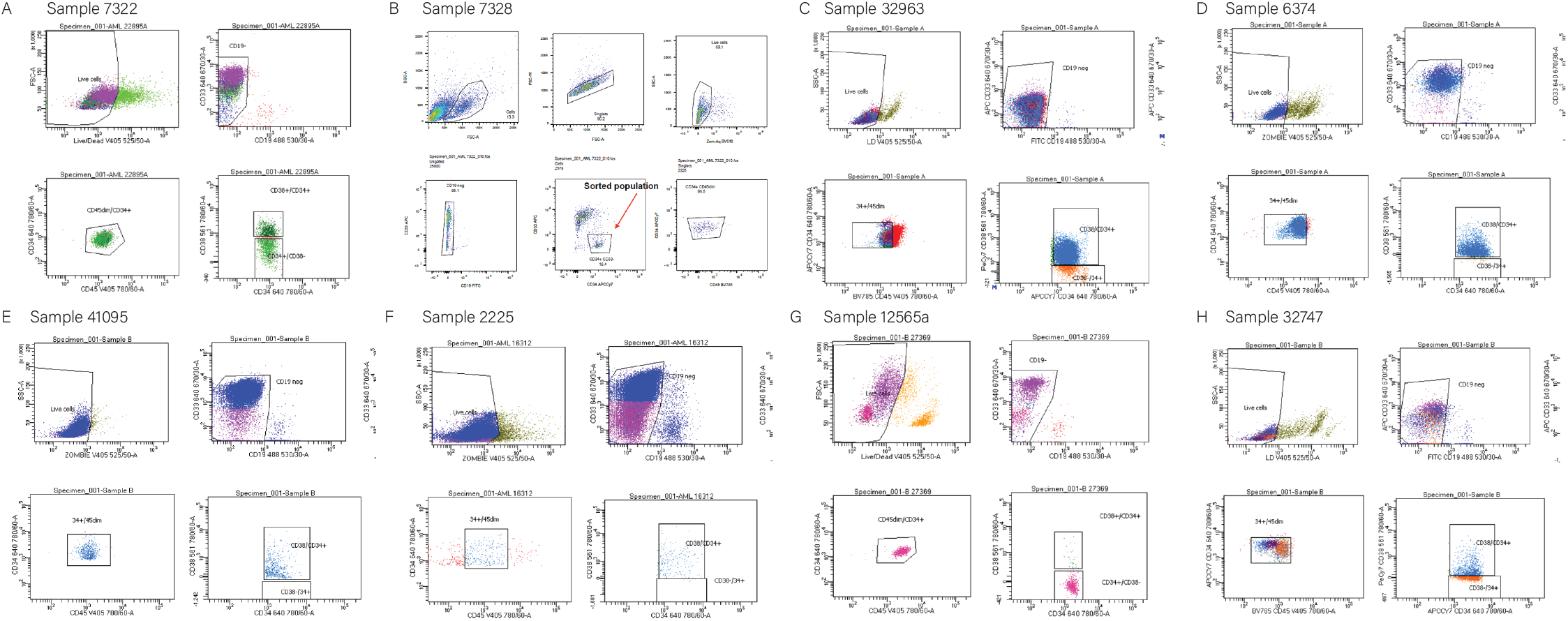
Examples of fluorescence-activated cell sorting gates.

## Supplementary Note 2: Inferring evolutionary parameters from clone size vectors using the tree similarity metric

### Stochastic simulation of ‘virtual AMLs’

We simulated the process of mutation acquisition and clonal expansion before AML onset using a multi-type branching process. Stem cells divide symmetrically at rate 1*/τ* = 1 per year and acquire driver mutations stochastically at a rate of *µ* per cell per year. Every driver mutation confers the same selective advantage *s*, depending on the selection level within the stem cell population, to the lineage and their effects combine additively, i.e. lineages harbouring two driver mutations are twice as fit as lineages harbouring only one driver mutation. This biases cell fate towards symmetric renewal and such cells produce 1 + *s* offspring on average in each generation. Clonal competition was modelled by allowing lineages to grow stochastically at a fitness relative to a growing mean fitness as the number of fit lineages increases. This maintains the total HSC population at a relatively constant size of *N* = 10^5 8,9^. Stochasticity of mutations and clonal expansions was modelled using Poisson processes. Cancer onset is defined as the event where any lineage acquires *k* driver mutations.

Using this computational model we performed simulations to generate 4000 3rd-hit events per *s* across *s* = 0%, *s* = 3%, *s* = 6%, *s* = 9%, *s* = 12%, *s* = 15%, *s* = 18%, *s* = 21%, *s* = 24%, *s* = 27%, *s* = 30% per year and 4000 4th-hit events per *s* across *s* = 9%, *s* = 12%, *s* = 15%, *s* = 18%, *s* = 21%, *s* = 24%, *s* = 27%, *s* = 30% per year, all at *µ* = 10^−5^ per year. Values of *s* below 9% per year produce implausibly low AML occurrence rates for *k* = 4 so were not considered except for the analysis on individual variation where we examined 1000 ‘virtual AMLs’ per *s* down to *s* = 6% per year. The clonal frequencies of clones ancestral to the *k* − *th* mutant at the time of ‘cancer onset’ were recorded for each event and stored as the corresponding clone size vector ***n*** = (*n*_0_, *n*_1_, …, *n*_*k*−1_).

### Tree similarity and set similarity

We extracted the clone size vectors for all the unambiguous preleukemic phylogenies (6 three-hit and 6 four-hit observed preleukemic trees) constructed from our single-cell experiments using the clonal frequencies of the main branches. We measured the set similarity of the 12 observed clone size vectors by summing the tree similarity of each observed clone size vector with its best-matched virtual AML tree. The set simlarities displayed in Figure 2 and 3 were normalized to show similarity calibrated to 1 for a perfect match i.e. all pairs of clone size vectors are completely identical. The tree similarity between any two clone size vector ***n*** and ***n***^*′*^ is calculated as 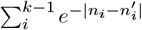. This metric is less sensitive to outliers than using the exponentiation of euclidean distance. We compared real, observed trees (***n***, purple lines) from our single-cell experiments with simulated ‘virtual AMLs’ (***n***^*′*^, pink lines) and found that the regime at *s* = 12% and *µ* = 10^−5^ shows the highest total similarity to the set of 6 three-hit and 6 four-hit trees from our single-cell experiments (Supplementary figure S6).

### Validation using published single-cell data

Morita et al. ^27^ reported the clonal architecture and mutational histories of 123 acute myeloid leukemia (AML) patients reconstructed from single-cell data. Of the 123 patients, 98 patients were analyzed for the single-timepoint sample collected at pre-treatment (*N* = 98). There were 93 (out of 123) *de novo* AML samples, among which 88 were not treated. These samples were sequenced on a narrower panel focusing on 19 AML genes. Based on additional published information on tree topologies from Schwede et al. ^55^, we extracted 44 three-hits (27 linear) and 23 four-hits (15 linear) clone size vectors. (Supplementary figure S7).

We compared the sets of *k* = 3 and *k* = 4 cases separately with ‘virtual AMLs’ simulated from the broader classes of parameters and found that *s* = 12% shows the highest similarity to the observed sets (Supplementary figure S8 and S9). This results holds and is not affected by how we divide the observed tree set into subsets (e.g. based on branching patterns, *k*…etc.). It remains highly consistent even when we reduce the number of simulated AMLs considered from each parameter regime (Supplementary figure S10). Trees from Morita et al. were directly inferred from bone marrow and peripheral blood mononuclear cells and in principle suffers less from subsampling biases. The highly consistent results with our inference despite low cell numbers in our experimental pipeline serves as a strong validation for our inference method.

**Fig. S6.**
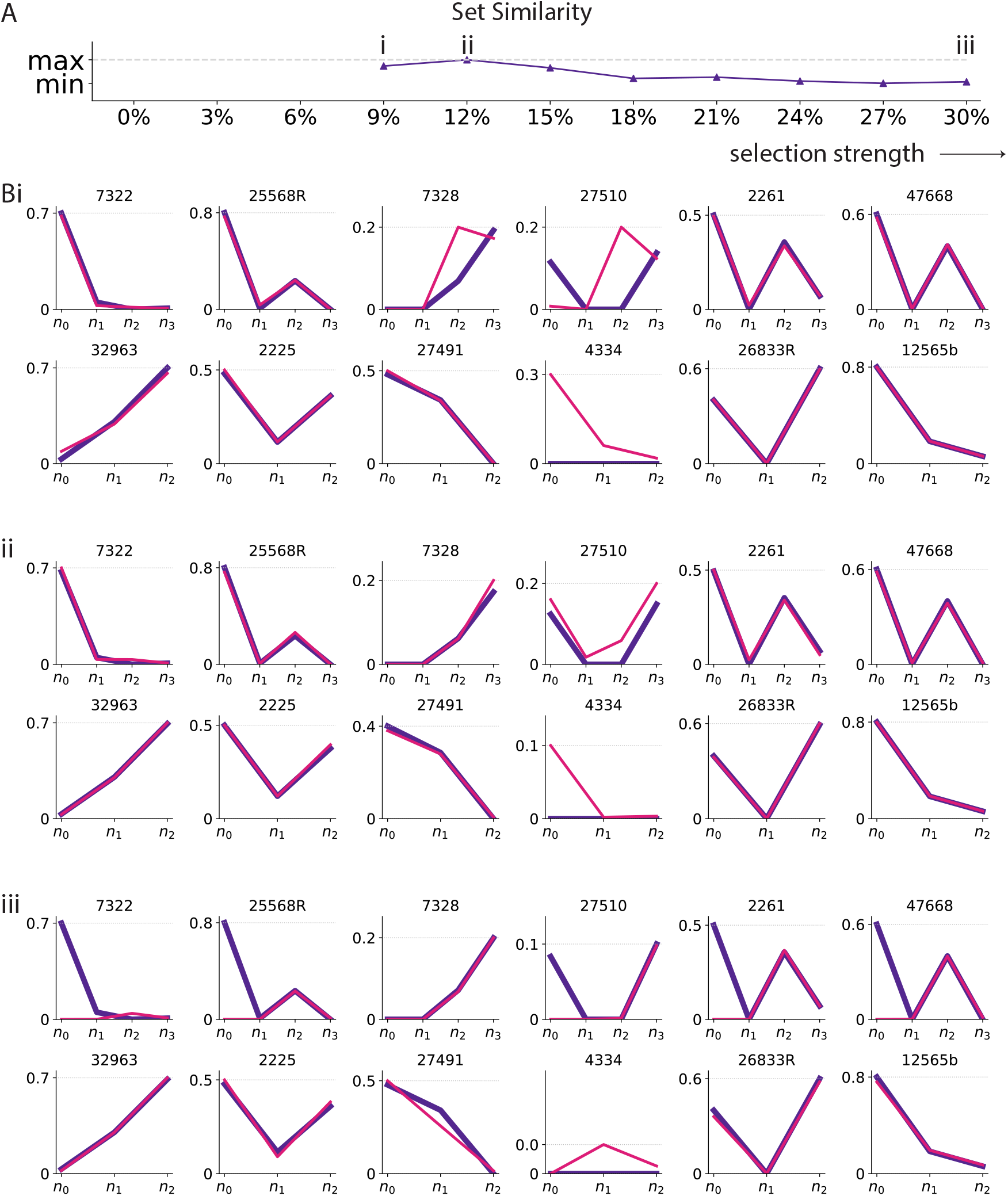
Set of all 16 unambiguous preleukemic trees shows highest similarity with ‘virtual AMLs’ at *s* = 12% per year. **(A)** Set similarity peaks at *s* = 12% per year. **(Bi)** The best-matched simulated tree (tree with highest similarity score, pink line) from 8000 virtual AMLs at *s* = 9% per year was chosen (with replacement) for each of the 6 three-hit and 6 four-hit preleukemic trees (purple lines) from our single-cell experiments. **(ii)** The best-matched simulated tree (tree with highest similarity score, pink line) from 8000 virtual AMLs at *s* = 12% per year was chosen (with replacement) for each of the 6 three-hit and 6 four-hit preleukemic trees (purple lines) from our single-cell experiments. The regime at *s* = 12% harbours the highest set similarity with our set of unambiguous preleukemic trees. **(iii)** The best-matched simulated tree (tree with highest similarity score, pink line) from 8000 virtual AMLs at *s* = 30% per year was chosen (with replacement) for each of the 6 three-hit and 6 four-hit preleukemic trees (purple lines) from our single-cell experiments.

**Fig. S7.**
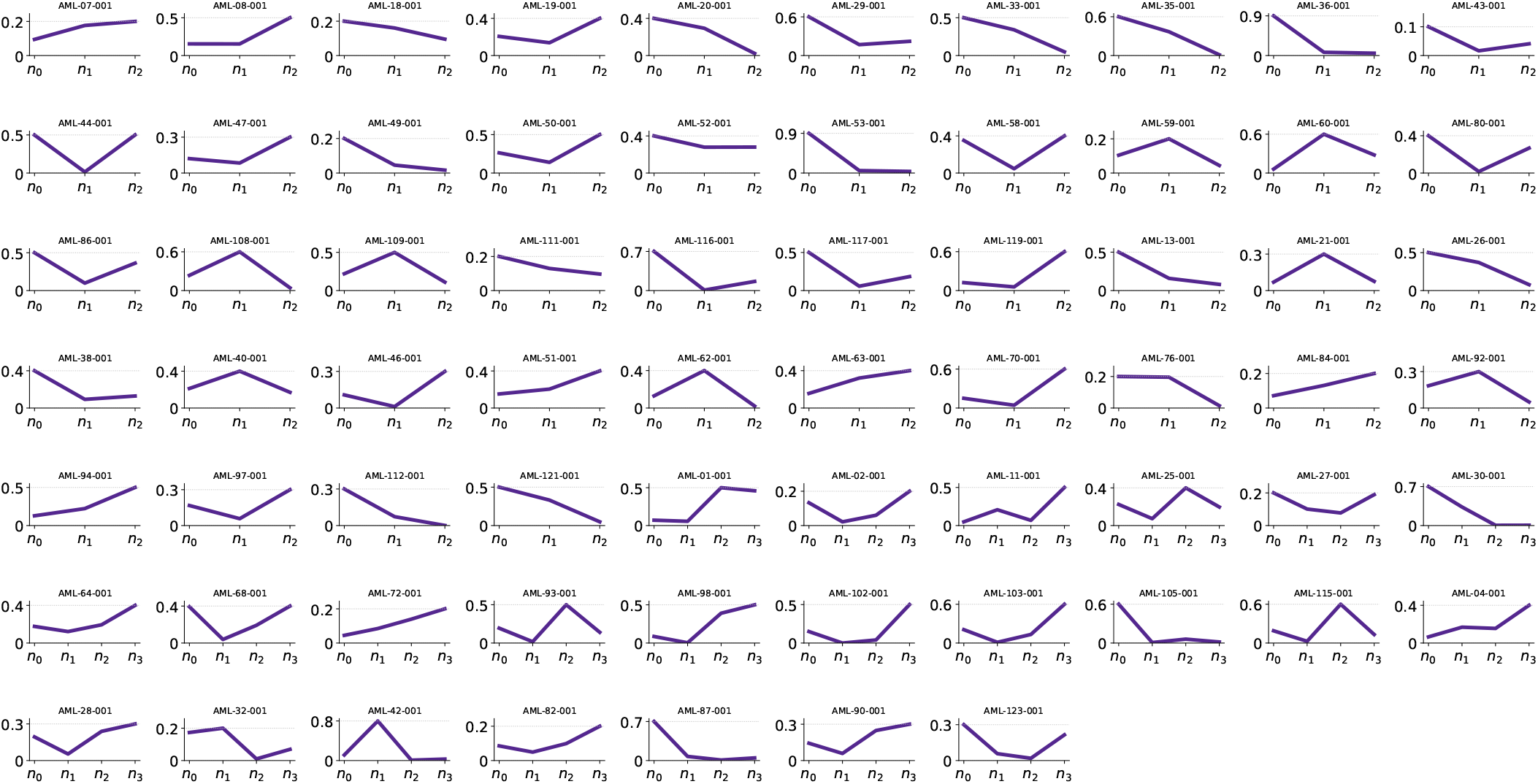
Sixty-seven clone size vectors were extracted from Morita et al. ^27^. The clonal frequencies of main branch clones were inferred from the published mutation order ^55^ and published psuedo-bulk mutation VAF differences (Morita supplementary table ^27^). For branched cases, side branch mutations were treated like they were not genotyped. Note that the published mutation orders were not always in descending order of published VAFs, indicating inherent uncertainties in both tree inferences and pseudo-bulk VAF measurements.

**Fig. S8.**
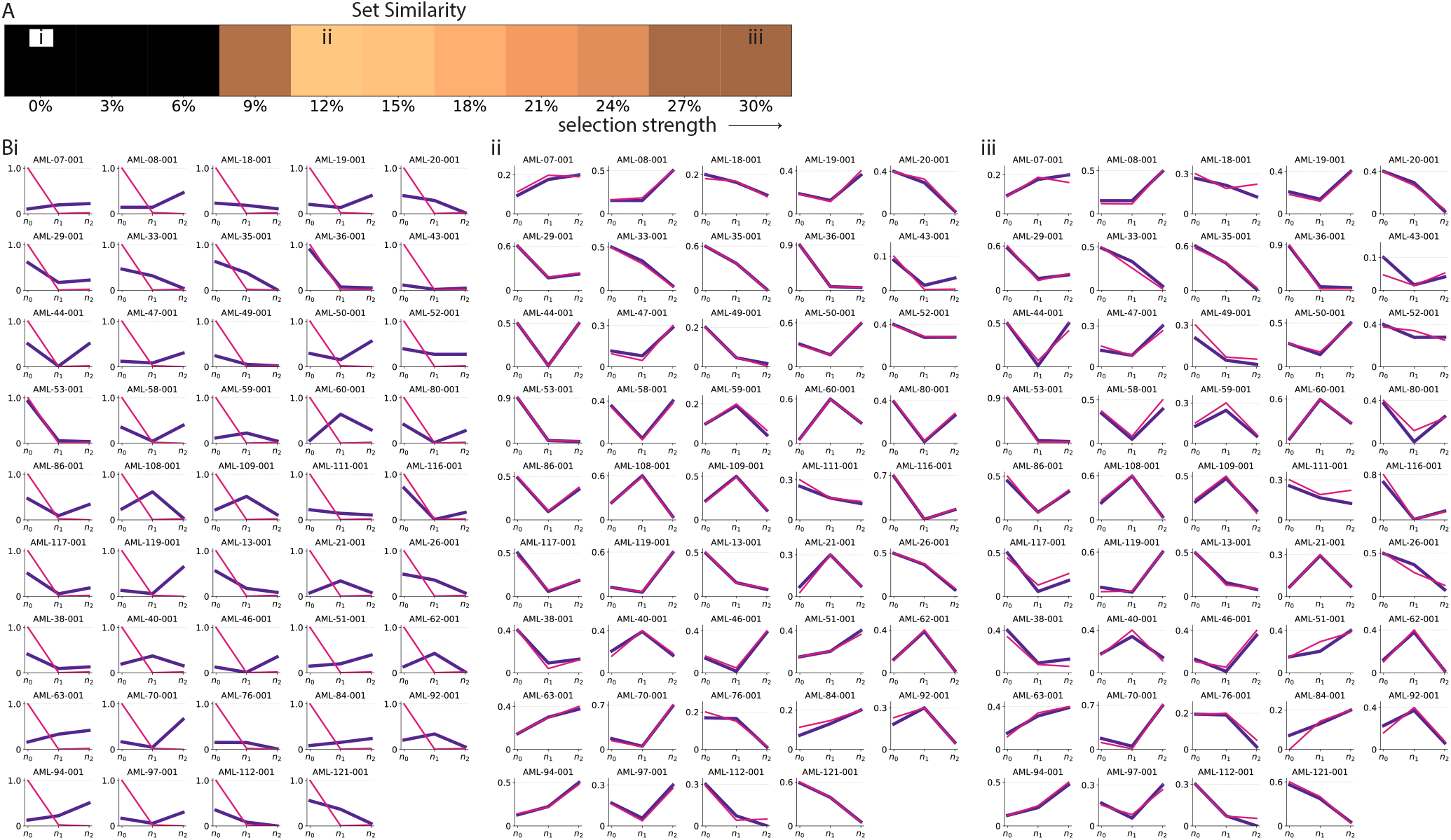
Moderate selection produces the most similar set of trees to Morita et al. ^**27**^ (*k* = 3). Considering 4000 trees simulated from every grid point of the parameter space, the most similar trees were identified for the 44 three-hit Morita cases. The similarity is summed across the entire set and reflected by the set similarity score. **(A)** Set Similarity score across the parameter space by comparing with all 44 three-hit cases. **(Bi-iv)** The most similar simulated tree (pink) from the corresponding parameter space is plotted against the real tree (purple line) for all 44 three-hit Morita trees.

**Fig. S9.**
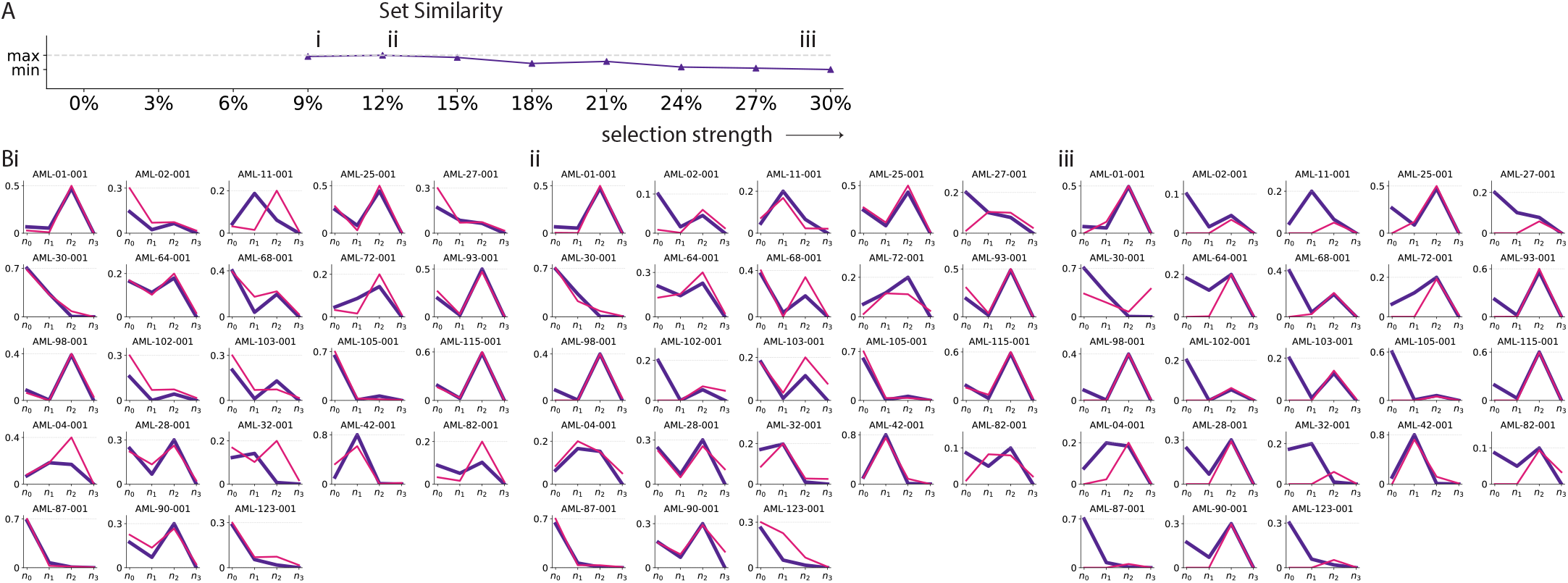
Moderate selection produces the most similar set of trees to Morita et al. ^27^ (*k* = 4). Considering 4000 trees simulated from every grid point of the parameter space, the most similar trees were identified for the 23 four-hit Morita cases. The similarity is summed across the entire set and reflected by the set similarity score. **(A)** Set similarity score across the parameter space by comparing with all 23 four-hit cases. **(bi-iv)** The most similar simulated tree (pink) from the corresponding parameter space is plotted against the real tree (purple line) for all 23 four-hit Morita trees.

**Fig. S10.**
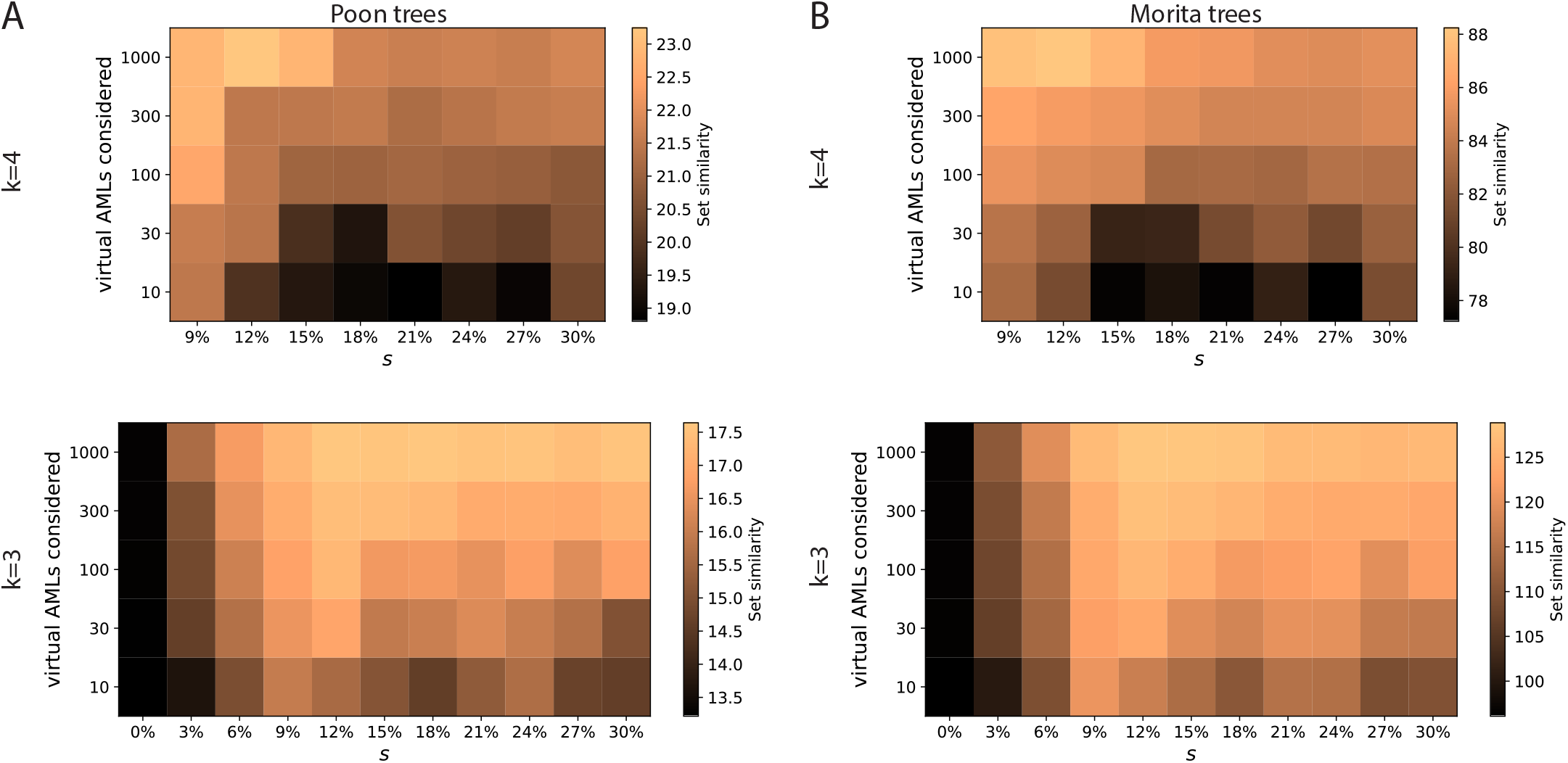
Tree compairson with Morita et al. in the neighbourhood of moderate selection *s* = 12%. Unsurprisingly, as the number of ‘virtual AMLs’ from each grid point considered is reduced, the similarity of the best-matched trees are also reduced and inference of the best-matched regime is noisier. However, *s* = 12% is largely the regime that produces the best-matched ‘virtual AMLs’.

## Supplementary Note 3: Inferring evolutionary parameters for individual subsampled trees using machine learning

### Extracting clone size vectors from stochastic simulations to account for subsampling

To evaluate the impact of subsampling effects we performed the same simulations as described in Supplementary note 2 to generate hundreds-of-thousands of virtual AMLs. However, to construct the clone size vectors this time we consider the absolute cell number that belongs to each clonal genotype after multinomial subsampling, instead of using clonal frequencies. Each virtual AML was subsampled *n* times and the probability of subsampling from a certain clonal genotype is equal to the cell fraction that clone constitutes in the total stem cell pool. The entries of a clone size vector therefore sum to *n* minus the number of subsampled side branch cells. We recorded 5000 3rd-hit and 1000 4th-hit events for each pair of (*µ, s*) values in the parameter space that produces cancer onset incidence within reasonable bounds: (10^−4^, 0%), (10^−4^, 15%), (10^−4^, 30%), (10^−5^, 15%), (10^−5^, 30%) for *k* = 4 and (10^−4^, 0%), (10^−4^, 15%), (10^−4^, 30%), (10^−5^, 0%),(10^−5^, 15%), (10^−5^, 30%),(10^−6^, 15%), (10^−6^, 30%) for *k* = 3 (Supplementary figure S11A).

**Fig. S11.**
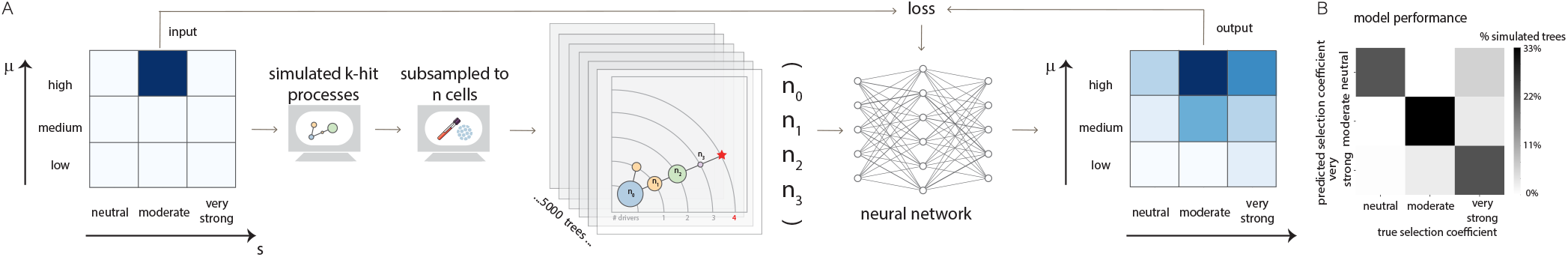
A statistical learning framework infers levels of selection and mutation encoded in preleukemic phylogenetic trees. (**A**) Stochastic simulations of the evolutionary dynamics were performed with known parameters for mutation rate *µ* and fitness effect *s* in a constant population of HSCs. For each simulation the phylogenetic tree was recorded and sub-sampled to *n* cells (*n* = 30, 100, 300, 1000) to model the effects of subsampling during the process of single-cell HSC sequencing. The resulting clone size vectors were then used to train a neural network classifier to infer the correct levels of selection and mutation. The classifier was trained on two-thirds of all simulated events and tested on the one-third of events that was held out. (**B**) We trained the models to classify across a range of selection strengths : neutral (*s* = 0% per year), moderate (*s* = 15% per year), and very strong (*s* = 30% per year). The performance of the classifier model in recognising selection strengths trained for three-hit processes after subsampling to *n* = 1000 is illustrated as an example. This particular classifier model achieves *>* 50% accuracy in assigning events to the correct class among 8 classes spanning across different selection strengths and mutation rates.

**Fig. S12.**
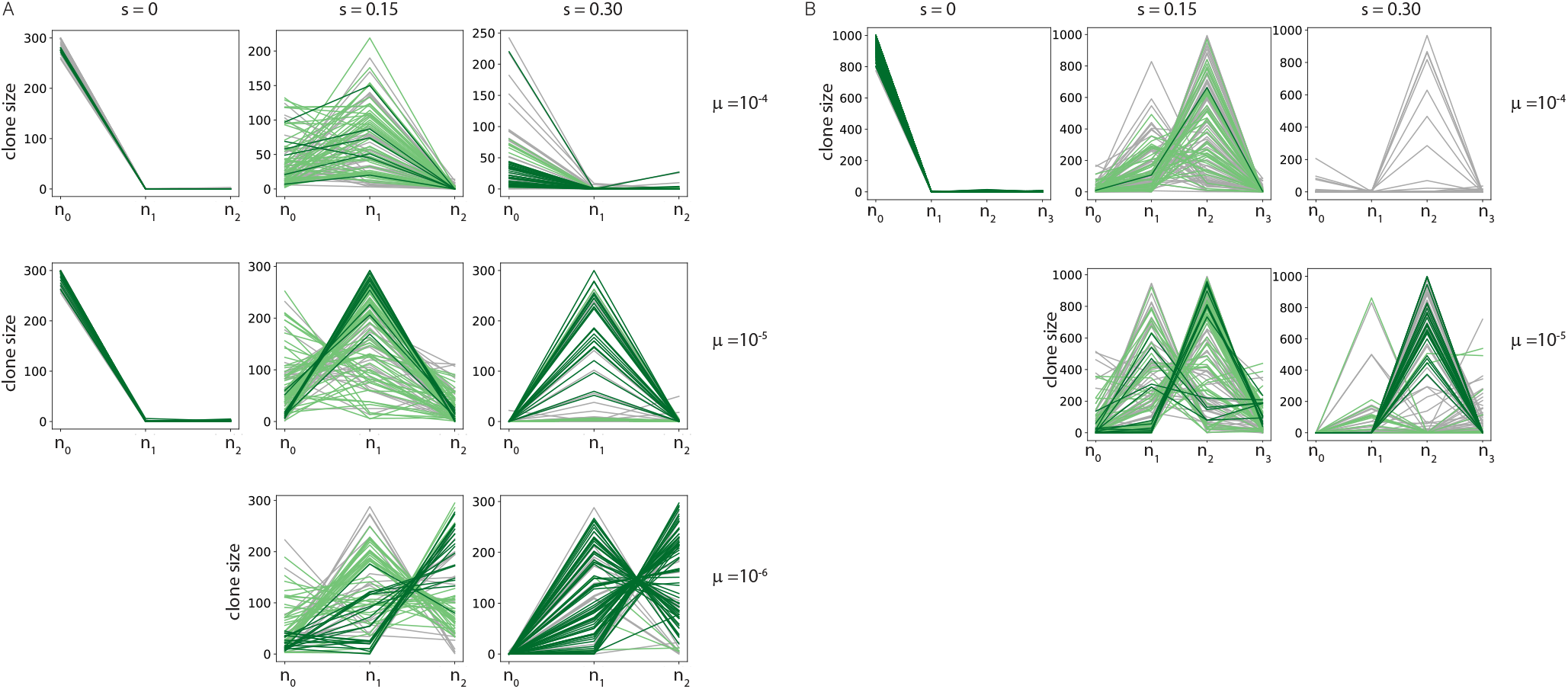
Visualization of clone size vectors belonging to the training set. **(A)** One example set of clone size vectors from subsampling ‘virtual AMLs’ (*k* = 3) down to *n* = 300. **(B)** One example set of clone size vectors from subsampling ‘virtual AMLs’ (*k* = 4) down to *n* = 1000. Models after training correctly recover the ground-truth i.e. correct (*µ, s*) with full consensus (10 out of 10) for clone size vectors highlighted in dark green and 80% consensus for those highlighted in light green.

### ML model for parameter inference

The ‘virtual AMLs’ were split into training (3335 (*k* = 4) events / 26680 (*k* = 3) events) and testing (1665 (*k* = 4) events/ 13320 (*k* = 3) events) datasets. Each dataset was subsampled ten times to create ten subsampled datasets (Supplementary Figure S12). We trained a fully connected neural network with 4 hidden layers for each subsampled dataset to classify the clone size statistics vector ***n*** (with corresponding *k* and *n*) by the parameter regime it was generated from. The models were trained using stochastic gradient descent and cross-entropy loss at a learning rate of 10^−8^ in batch sizes of 32 over a number of epochs. The number of epochs was decided based on the typical convergence in test accuracy across models trained using different seeds. Model accuracies converge to ≈ 65% (*n* = 1000), ≈ 62% (*n* = 30), ≈ 52% (*n* = 10) for *k* = 3 and ≈ 60% (*n* = 300), ≈ 60% (*n* = 30), ≈ 36% (*n* = 10) for *k* = 4 (Supplementary Figure S13). Down-sampling reduces the signal-to-noise ratios and typically results in lower classification accuracies. However models for *n* ≥ 30 can still achieve test accuracy beyond 50% (Supplementary figure S11B). By training on this large number of virtual cancers the models were able to learn how patterns in the phylogenetic trees encoded the selection strengths and mutation rates that were operating during preleukamic evolution.

We applied the fully trained neutral network to each of the 16 experimental trees to infer the patient-specific levels of selection and mutation which were operating during preleukemic evolution. For each real patient pHSC tree, we applied 10 models trained on independently sampled datasets to infer its parameter regime based on the corresponding number of hits *k* and the closest *n* to its sample size (Supplementary Figure S14). The number of models producing a certain inference on the parameter space was counted and the sum across the ten models represents the classification outcome of the sample (Supplementary Figure S15). This ensemble approach provides an estimate of variance for the classifier in its predictions.

The following real trees were classified with the (*n* = 1000, *k* = 4) ensemble classifier: ID7322, ID7328, and ID25568R. Tree ID2225 and ID32963 were classified with the (*n* = 300, *k* = 3) ensemble classifier and tree ID27491 was classified using the (*n* = 100, *k* = 3) ensemble classifier. For trees with lowest cell output, tree ID4334 was classified with the (*n* = 30, *k* = 4) ensemble classifier and trees ID27510 and ID2261 were classified using (*n* = 30, *k* = 4) ensemble classifier. The smallest ambiguous trees ID12565b, ID26833R and ID47668 was fed to the (*n* = 10, *k* = 3) and (*n* = 10, *k* = 4) ensemble classifier respectively. The remaining patient trees were not applied to our models as clonal genotype information was insufficient.

The classification outcomes applied to observed preleukemic trees indicate that classifiers trained on independently subsampled datasets typically converge on the inferred selection levels (Supplementary figure S15). The inferred selection levels are largely consistent with our previous inferences which indicates that biases introduced by subsampling only influence a small subset of the observed trees: 27510 and 26833 (and possibly ID25568R, 2261, 47668 to a lesser extent).

**Fig. S13.**
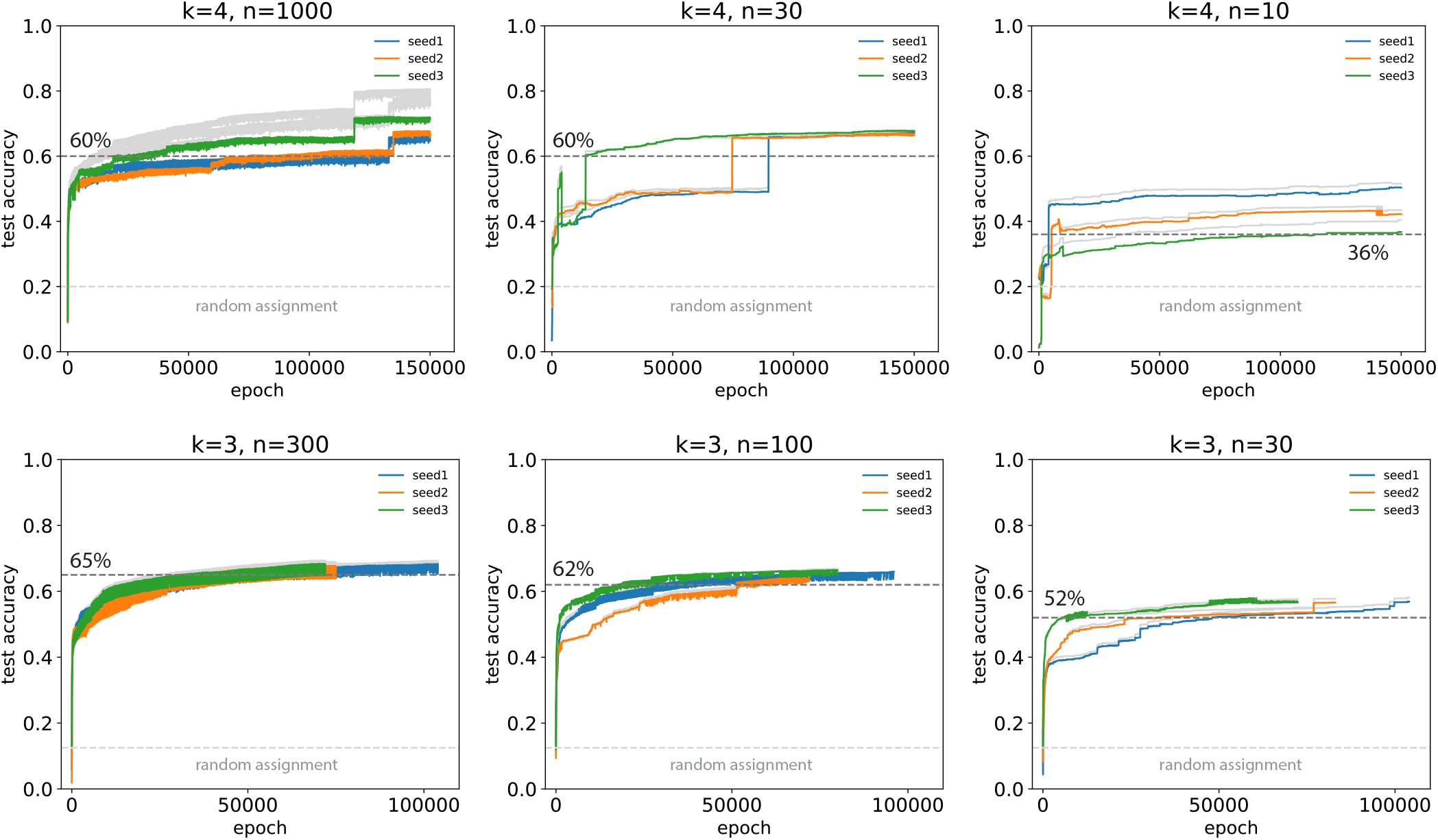
Typical test accuracies after training. Using one single training/testing set of clone size vectors for each (*k, n*), we trained models using different random seeds to show that the models typically converge to the labelled test accuracies. All classifier models used to classify real patients pHSC trees had been subsequently trained until they reach test accuracies above or similar to these labelled accuracies. The expected test accuracies based on random assignment are shown for comparison. Training accuracies are also plotted as grey lines for each seed which shows that there is no over-fitting.

**Fig. S14.**
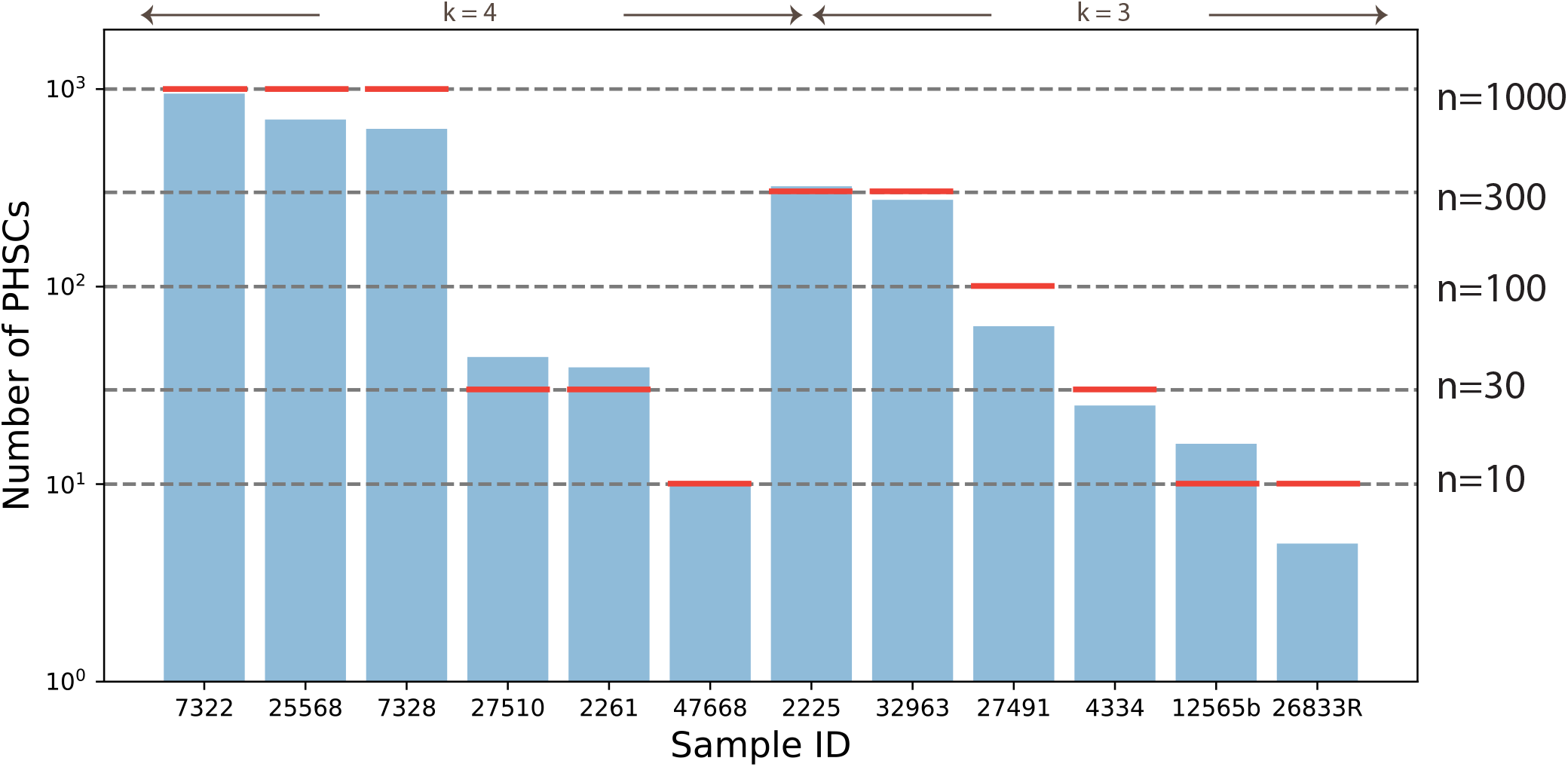
Choice of *n*. Each sample was matched to the closest *n* in logarithmic scale (red line) based on the number of preleukemic HSCs yielded after single-cell DNA sequencing. The number of driver mutations determines *k*. All twelve samples with unambiguous pHSC trees fed into the machine learning framework are shown.

**Fig. S15.**
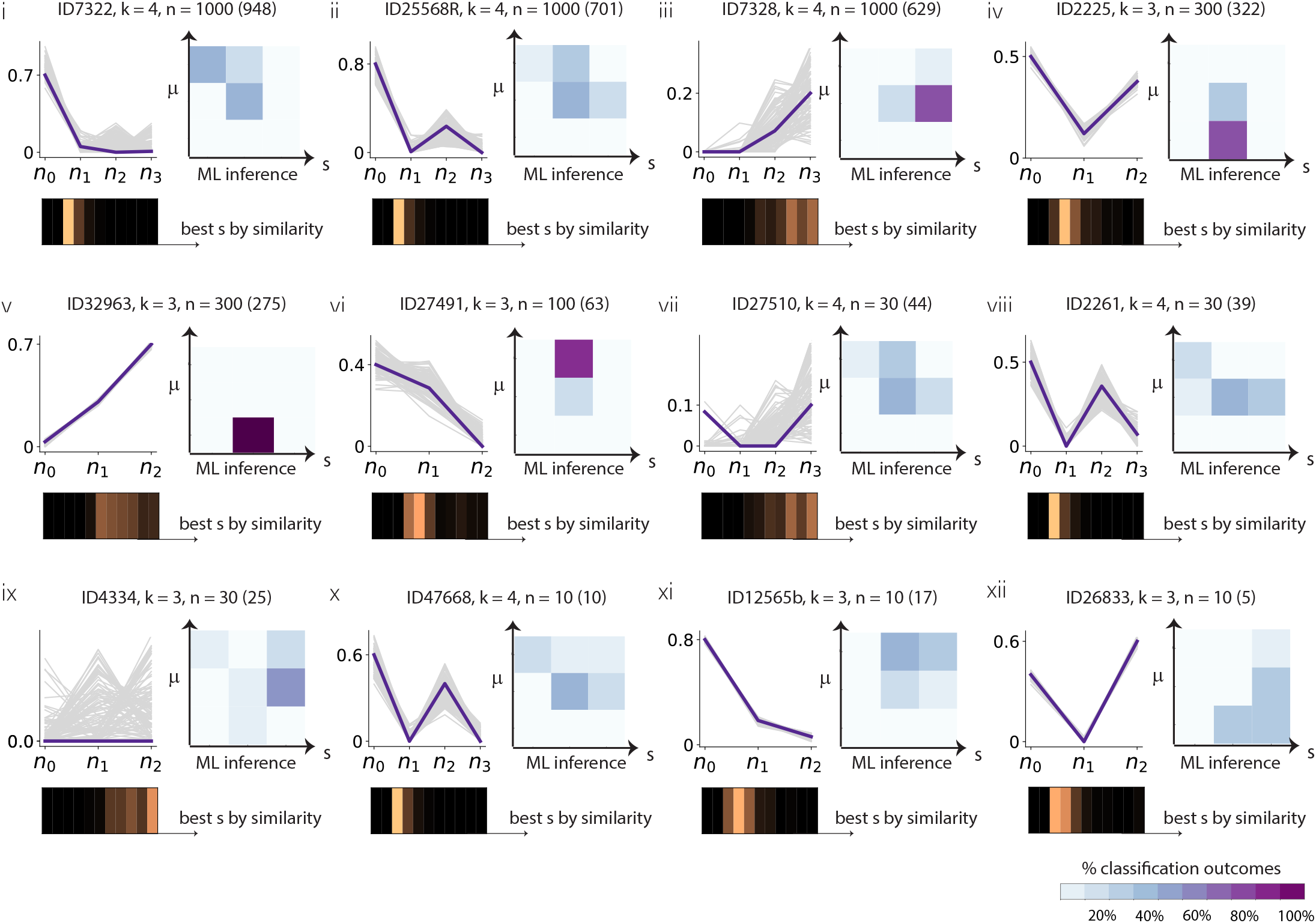
Classifier predictions for patient preleukemic HSC trees. We applied our classification scheme for the twelve experimentally observed phylogenetic trees (6 four-hit trees and 6 three-hit trees) without unambiguous clonal genotypes. For each tree, ten classifier models of the relevant number of hits *k* and *n* closest to the true pHSC number (bracketed number) were applied to generate the final classification outcome (displayed on the right of the tree). It represents the fraction of classifiers producing a specific outcome on the the parameter space. We compared this outcome with our previous inference using direct clonal frequencies (displayed at the bottom of the tree). The inferred selection levels are largely consistent with our previous inferences which indicates that biases introduced by subsampling only influence a small subset of the observed trees: 27510 and 26833 (and possibly ID25568R, 2261, 47668 to a lesser extent).

**Fig. S16.**
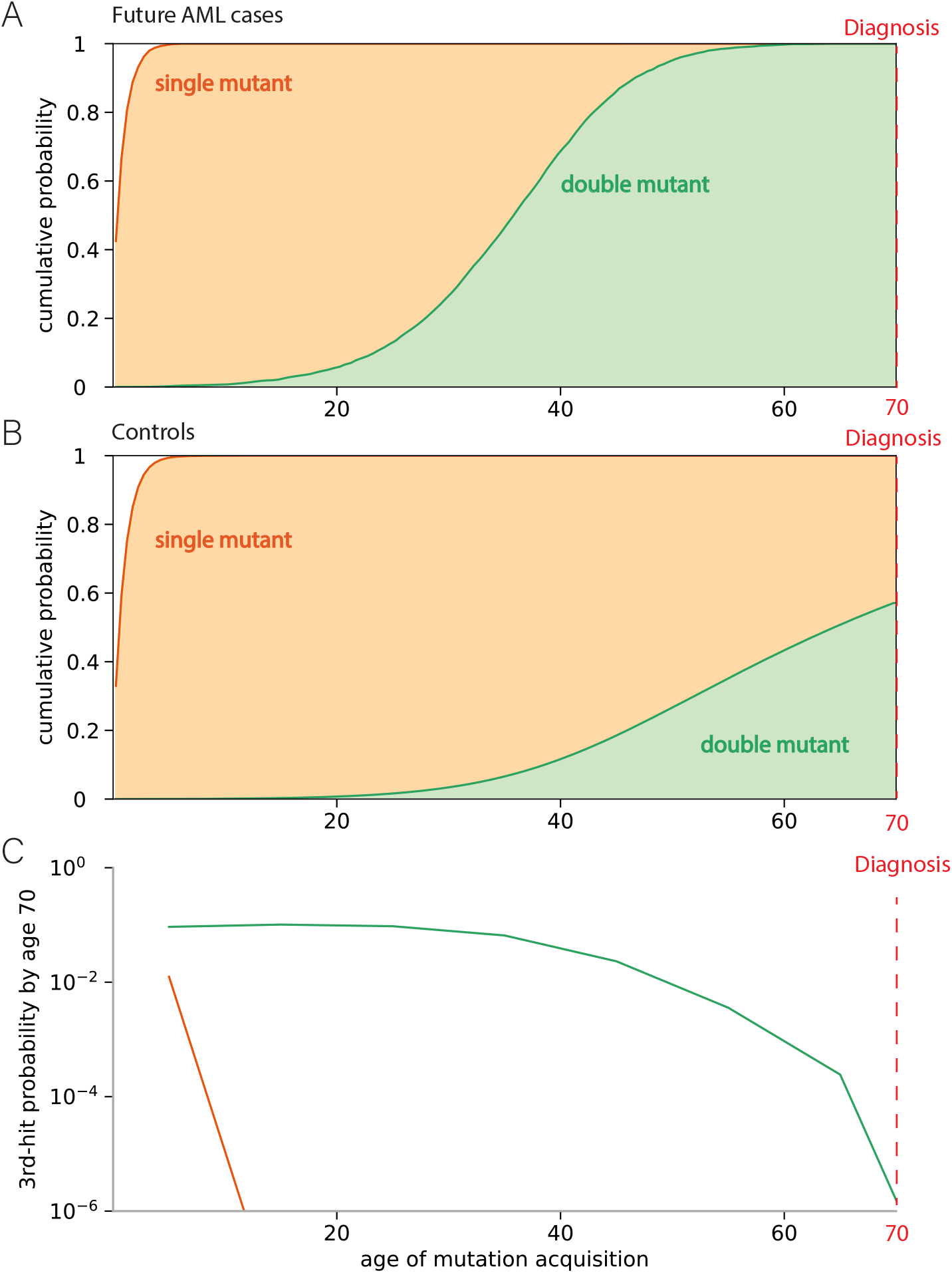
Early double-mutant clones identify a high-risk group for AML transformation. **(A)** Distribution of mutation acquisition times in simulated individuals (under moderate selection and at medium mutation level) who acquired 3rd-hit by age 70. **(B)** Distribution of mutation acquisition times in unselected simulated individuals, showing marked differences compared to simulated individuals destined to develop AML by age 70. **(C)** The absolute risk of ‘virtual AML’ (three-hits) transformation before age 70 is dependent on when single (orange) and double (green) mutant clones are acquired.

**Fig. S17.**
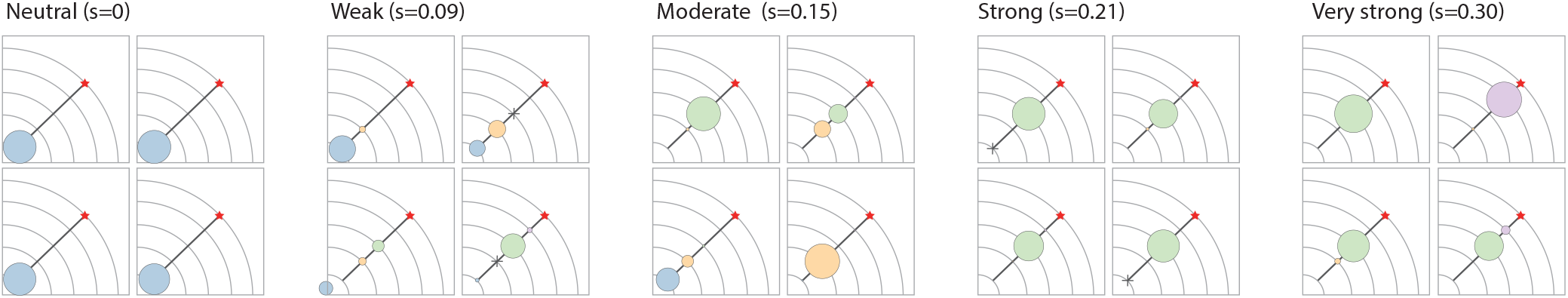
Tree pattern differences. ‘Virtual AMLs’ simulated over a range of selection strengths: *s* = 0.00, *s* = 0.09, *s* = 0.15, *s* = 0.21, *s* = 0.30 show visible differences.

**Fig. S18.**
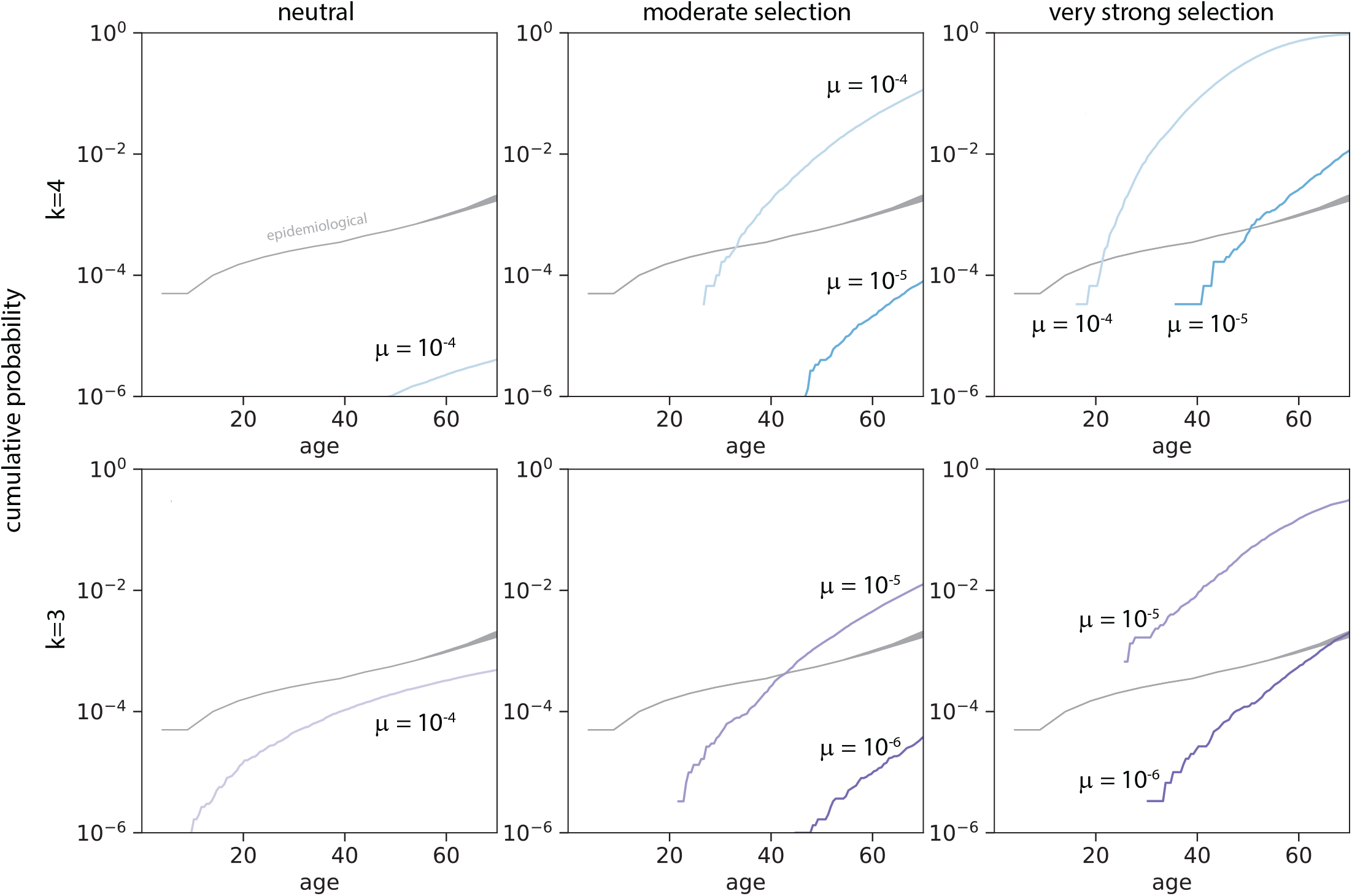
Cancer onset rates based on *k*-th staircase model compared to epidemiological rates. The *k*-th hit staircase model predicts cumulative rates of ‘virtual AMLs’ across time. For decreasing selection strengths, the corresponding mutation rates needed to produce reasonable cumulative incidence rates increases. Predicted cancer onset rates for *k* = 3 and *k* = 4 (neutral: *s* = 0%, moderate selection: *s* = 15%, strong selection: *s* = 30%) and are shown alongside observed epidemiological data.

## Notes

### Competing Interest Statement

The authors have declared no competing interest.

### Summary of Updates

This version of manuscript has been revised to include 67 additional phylogenetic trees from AML patients of an independent single-cell data.

